# Conditional Spatial Classification of Expert-Confirmed Interictal Epileptiform Discharge Epochs: An EEG-ECG Ablation and SHAP Analysis

**DOI:** 10.64898/2026.08.13.744348

**Authors:** Al Mukshit Plabon, Abdul Mukit, Md. Neyamul, Omar Faruk Jehady, Fatima Tuz Zuba, Md. Faisal Mina, Torikul Islam

## Abstract

Interictal epileptiform discharges (IEDs) are diagnostically important EEG abnormalities observed between seizures. This study addresses a conditional spatial-classification task where every analyzed four-second epoch had already been reviewed and confirmed by experts as containing an IED, and the model assigned that epoch to one of five predefined scalp-distribution categories (generalized, frontal, temporal, occipital, or centro-parietal). The analysis therefore does not evaluate IED-versus-non-IED detection. After preprocessing, 2,514 IED-labelled epochs were analyzed using identical stratified epoch-level partitions, SMOTE based training, 26 handcrafted features per included channel, and multiple machine-learning classifiers. A staged channel ablation compared 19-channel scalp EEG, 21-channel EEG with ECG, and the complete 29-channel input containing scalp EEG, referential, ECG, and EMG channels. The best EEG-only result was obtained with linear discriminant analysis (88.89% test accuracy). CatBoost achieved 93.25% on EEG with ECG channel and 94.44% with the whole channel set. All eight directly comparable classifiers showed numerically higher test accuracy after ECG channel was added; for CatBoost, the increase was 6.35 percentage points. In the EEG with ECG channel, CatBoost model on ECG channel on right and left arm received respectively 15.79% and 15.12% of normalized global SHAP attribution, and beta-band power was the leading of all features (18.76%). These SHAP values indicate model-specific predictive contributions and do not establish physiological biomarkers, causal autonomic mechanisms, or clinical localization. The findings support a limited methodological conclusion which is ECG-derived features were associated with improved internal epoch-level categorization of expert-confirmed IED epochs. They do not establish IED detection, artifact rejection, independent EMG effects, or generalization to unseen patients.

## Introduction

Epilepsy is a disorder of the nervous system in which seizures occur spontaneously and without any triggering factors due to abnormal functioning of the neurons [1], [2]. Electroencephalogram (EEG) plays a pivotal role in clinical assessment by capturing the electrical activity of the brain and potentially identifying interictal epileptiform discharges (IEDs) between epileptic seizures [3], [4], thus helping to detect the presence of abnormalities. The morphology and distribution of the IEDs can help the electroclinical interpretation of the seizure, but the scalp category of the IED does not define the source localization, the seizure onset zone or the epileptogenic zone.

Based on the distribution in the scalp, IEDs can be classified as generalized or focal. The generalized IEDs were classified as one class, and the focal IEDs were divided into frontal, temporal, occipital and centro-parietal classes[5]. Importantly, each analyzed time frame had already been determined to be a time of IED activity. The task thus involved conditional spatial classification where the model predicted one of five predefined labels related to the scalp distribution of the IED. Throughout the article, the term "spatial classification" is used without the connotation of source reconstruction or identification of the seizure onset zone.

The problems of IED detection and conditional spatial classification are different problems [6], [7]. Detection involves determining if an unselected EEG segment contains an IED and comparing it to non-IED background, artifacts and IED-like transients[8]. The following analysis only starts after this decision is made by expert annotation. It therefore does not attempt to estimate detection sensitivity or specificity, false-positive burden or artifact rejection, or performance in continuous EEG review. It is limited to post-annotation categorization of IED epochs identified by experts.

While there has been significant focus on binary IED detection, there are limited studies on multi-class categorization following detection of an IED [5],[9]. This downstream task is more clinically focused than detection but offers a controlled environment for examining whether channel groups contain information that is useful for discriminating between a set of categories in the scalp-distribution task. It also allows to ask a new question (direct modality-ablation) which is not answerable using only feature-importance analysis[10].

The source recordings contain scalp EEG, referential, Electrocardiogram (ECG) and Electromyogram (EMG) signals[5]. Three channel configurations, 19-channel scalp EEG, scalp EEG with two ECG channels, and a full 29-channel input were compared. The EEG-versus-EEG with ECG comparison focuses on the numerical change that comes from combining the ECG components. The comparison between EEG with ECG and the full input was made with the joint addition of four referential and four EMG channels, and does not distinguish between the contribution of the EMG channels and the contribution of the referential channels.

We used SHapley Additive exPlanations (SHAP) to explain the variables in the channel features that led to each fitted model [11]. SHAP importance depends on the dataset, the features used, the model and the target [12]. It is not a physiological biomarker, it has not shown to be a marker of clinical validity, nor has it proven to find IEDs[13]. This precaution is particularly relevant for ECG and EMG variables, as they can contain patient identity, sleep-wake state, recording conditions, contamination, movement or other specific structure of the dataset[14], [15], [16].

The goal of the proposed work was to (1) assign expert-validated IED epochs to one of five pre-defined spatial categories; (2) compare different machine learning classifiers on this conditional task; (3) quantify the contribution of ECG features when added to scalp EEG; (4) compare performance of EEG with ECG versus complete 29-channel input; and (5) characterize the contribution of model-specific features and channels using SHAP without interpreting them as validated physiological biomarkers.

## Methodology

A conditional spatial-classification pipeline was used, based on the five classes of expert-confirmed IED epochs, for the study. The target classes were generalized, frontal, temporal, occipital and centro-parietal. Different channel configurations were tested using an identical modelling workflow and then the trained classifiers were analyzed using SHAP.

For all experiments, the set of experts confirmed epochs of IEDs were used to apply the same workflow as depicted in Fig 1. Twenty-six unique handcrafted features were extracted independently for each channel included. Three feature matrices were created: (1) 19 scalp EEG channels, (2) 19 scalp EEG channels and ECG RA, and (3) the 29 channels input, which includes 19 scalp EEG, 4 referential and 4 ECG channels. For each of the classifiers, a flat vector of channel features was provided. Coordinate of the electrodes, adjacency matrices, inter-electrode distance, topographic map and outputs of electrode source reconstruction were not used. The spatial information was only implicitly conveyed in the feature names of each channel.

### Proposed Methodology

**Fig 1.**
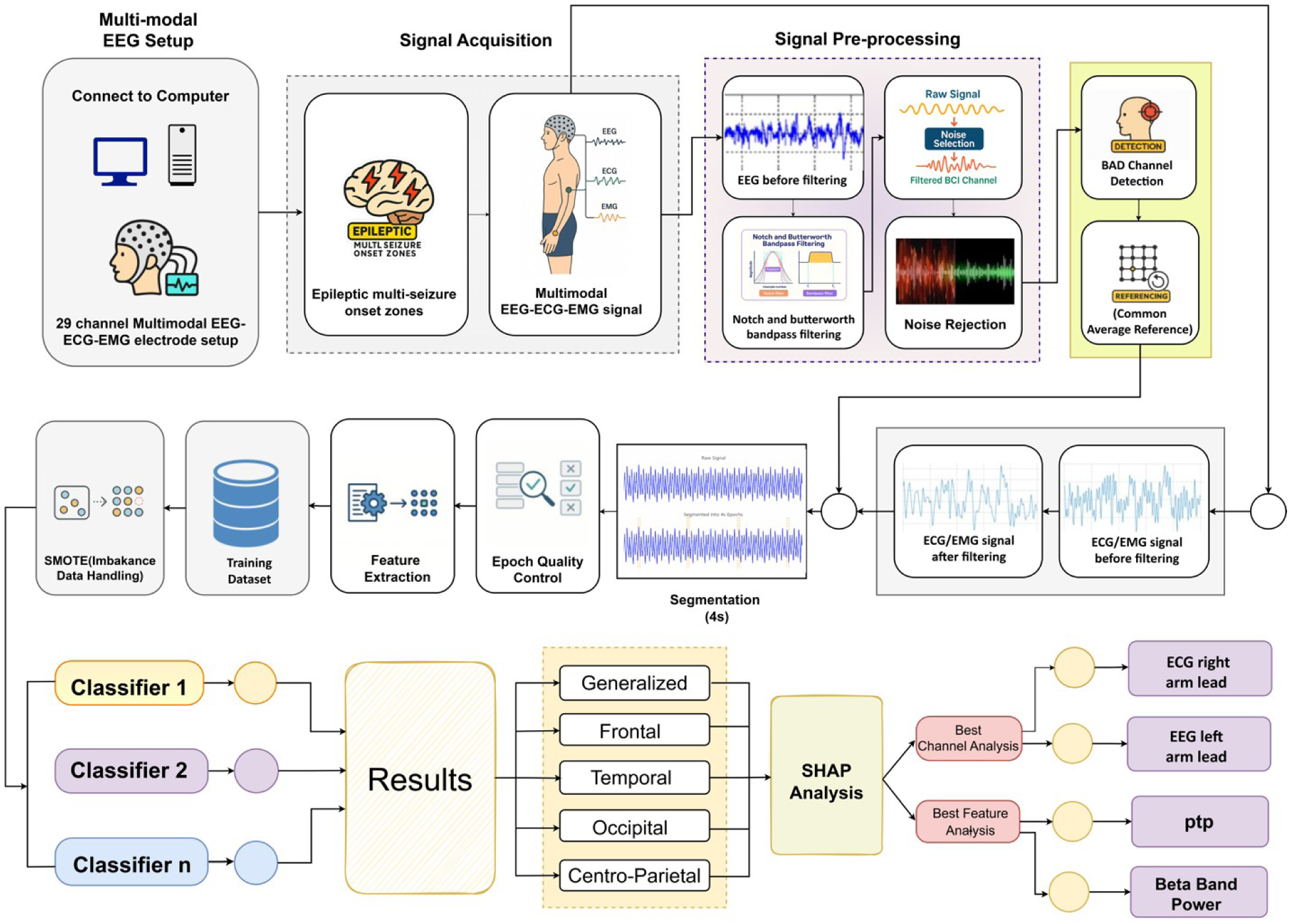
Study workflow for preprocessing, channel specific feature extraction, conditional five-class spatial classification, channel set ablation and SHAP analysis.

### Dataset

There are 29 channels on the recordings: 19 scalp EEG channels positioned using the international 10-20 system illustrated in Fig 2, four scalp referential channels (T1, T2, A1, and A2), 2 channels for ECG (RA and LA), and 4 channels for deltoid EMG (bilateral) depicted in Fig 3. Table 1 lists the channel groups and the location where they were recorded. The ECG RA and ECG LA are referred to as lead24 and lead25 in the EEG with ECG SHAP outputs, while this was the original feature naming convention.

**Table 1:** Recording of various signal types including the position of electrodes.

| **Type of recording electrodes** | **Number of Electrodes** | **Position of the Channel** |
| --- | --- | --- |
| EEG | 19 | Fp1, F3, C3, P3, O1, F7, T3, T5, Fz, Cz, Pz, Fp2, F4, C4, P4, O2, F8, T4, T6 |
| Referential | 4 | T1, T2, A1, A2 |
| ECG | 2 | Right Arm (RA) and Left Arm (LA) |
| EMG | 4 | Deltoid muscles: Two channels from the right deltoid region and two channels from the left deltoid region (bilateral). |

**Fig 2.**
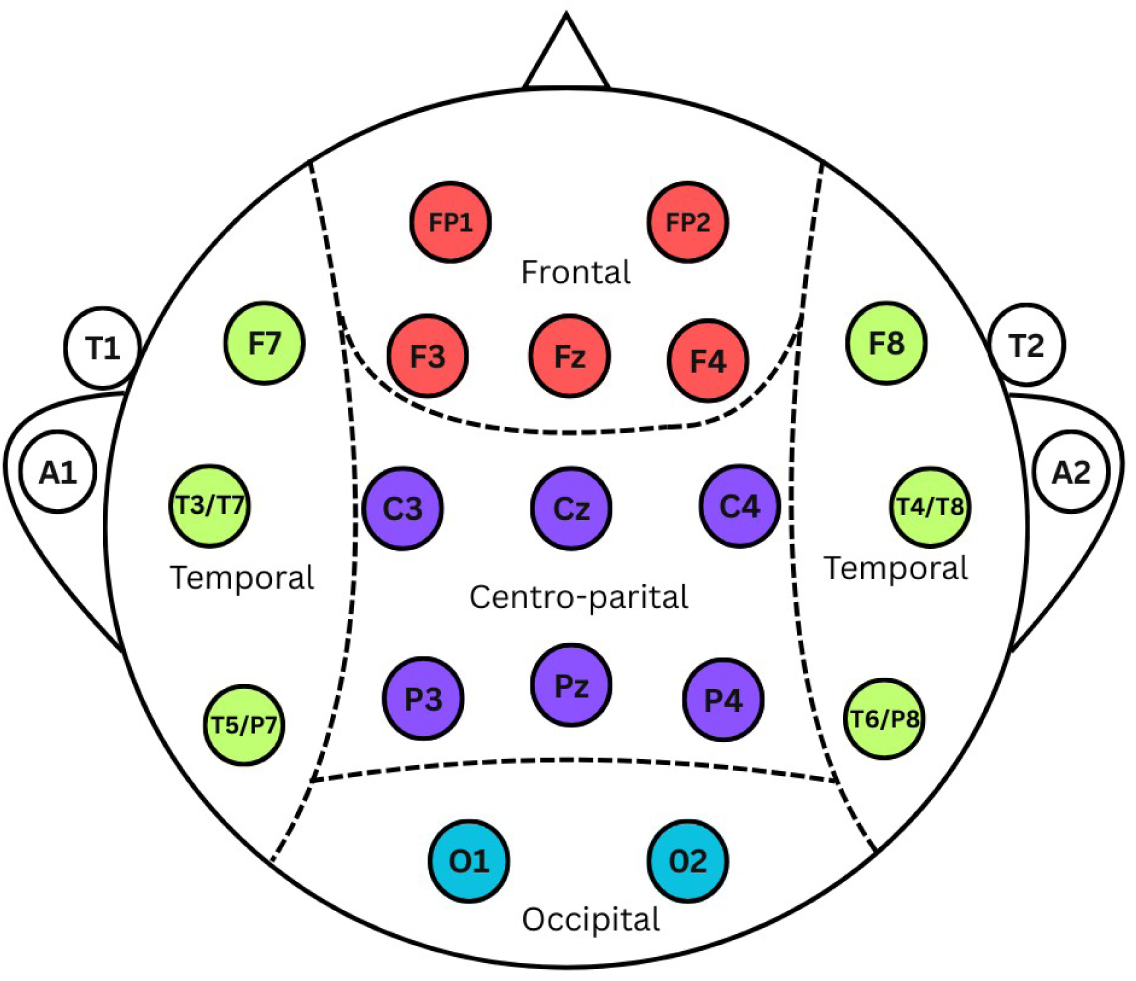
EEG electrode position and the predefined five spatial categories of IED. The "Fp" refers to frontopolar, “F” to frontal, “C” to central, “P” to parietal, “O” to occipital, and “T” to temporal locations.

**Fig 3.**
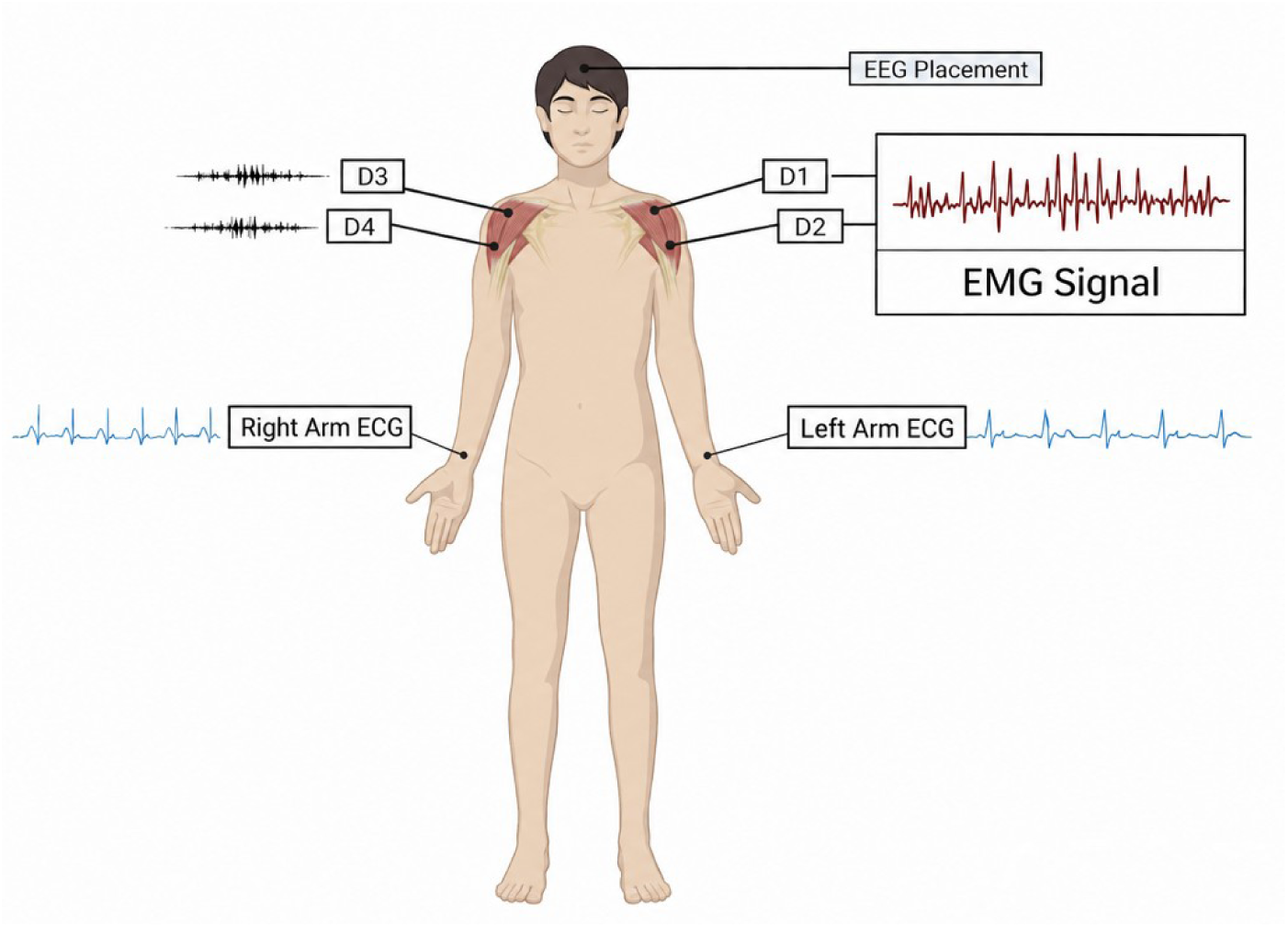
For an electrode arrangement in the ECG and EMG. The right and left arm ECGs are called RA and LA. Bilateral Deltoid EMG channels D1-D4 are indicated.

The open data set was obtained from the Epilepsy Center of Peking Union Medical College Hospital [5]. It has an audio content from 84 participants. Fifty-two recordings had at least one expert confirmed IED and 32 recordings were reported as normal EEG recordings. There were 20 minutes of data from each participant, which amounted to a total of 28 hours of data. There are 22,933 four second epochs in the source dataset that are not labeled as IEDs, and 2,516 four second epochs labeled as IEDs. During quality control, two IED epochs were discarded, resulting in 2,514 expert-confirmed IED epochs that were used for the present analysis. A class of non-IED epochs was not included as a target class.

MATLAB and NumPy formats of the recordings and the epoch labels. Table 2 lists the number of epochs of the original 2,516 epochs that were labeled with IEDs.

**Table 2:**
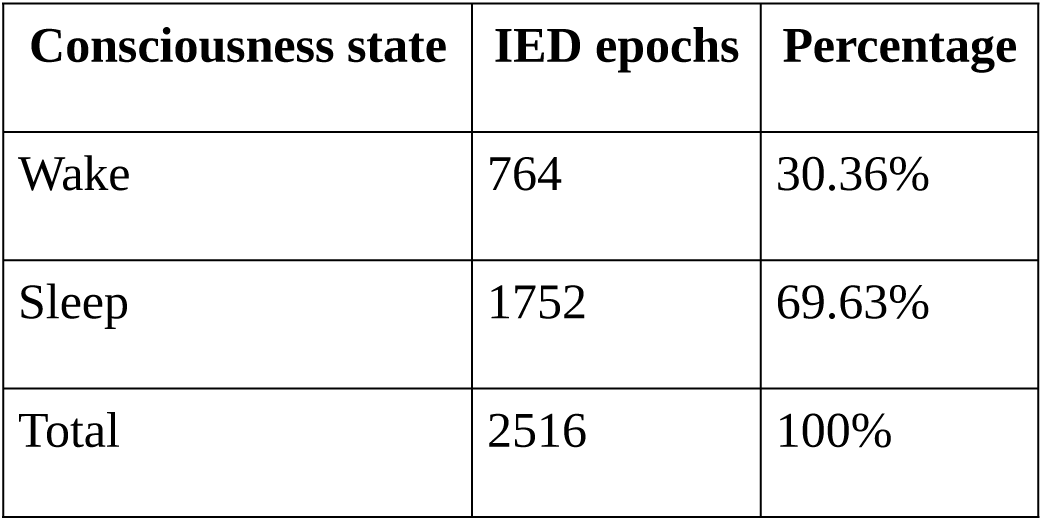
Consciousness state across IED epochs at 4 sec.

| Consciousness state | IED epochs | Percentage |
| --- | --- | --- |
| Wake | 764 | 30.36% |
| Sleep | 1752 | 69.63% |
| Total | 2516 | 100% |

### EEG Signal Preprocessing

The four-second epochs were linearly detrended, notch filtered at 50 Hz and at 100 Hz, and band-pass filtered between 0.5 and 40 Hz with a Butterworth filter [17]. Each channel configuration was pre-processed in the same manner prior to construction.

The composite criterion of robust log-variance outliers and low interchannel correlation was used to identify noisy or defective EEG channels. Detected bad EEG channels were interpolated from neighboring channels [18]. Independent component analysis was only used on the EEG channels to remove ocular artifacts while retaining the ECG and EMG signals [19]. All scalp EEG channels were then re-referenced to a common average.

Two epochs of wake state were rejected due to the peak-to-peak amplitude being above a strong median absolute deviation (MAD) threshold. The final dataset therefore contained 2,514 IED epochs: 762 wake epochs (30.31%) and 1,752 sleep epochs (69.69%).

### Feature Extraction

Each kept 4 second epoch was feature extracted. The 26 features were computed for each included channel, yielding 494 variables for EEG, 546 variables for EEG with ECG, and 754 variables for the 29 channels of input (EEG, ECG and EEG). Features were divided into three categories: time domain, frequency domain and nonlinear.

#### Time-domain Features

The variables used in time domain are mean, median, standard deviation, skewness, kurtosis, peak-to-peak amplitude, waveform length, slope sign changes, Hjorth mobility and Hjorth complexity.

#### Frequency-domain Features

Welch power spectral density estimates were used to compute frequency domain variables. Absolute band power was calculated for each band: delta (0.5-4 Hz), theta (4-8 Hz), alpha (8-13 Hz) and beta (13-30 Hz) as well as the ratio of each band power, peak frequency and median frequency.

#### Nonlinear Features

Nonlinear variables were approximate entropy, sample entropy, permutation entropy, spectral entropy, correlation dimension, detrended fluctuation analysis and the Hurst exponent [20].

### Data Splitting

The 2,514 epochs labeled with IEDs were split into epochs, with an 80:10:10 epoch-level split: 2,011 training epochs, 251 validation epochs, and 252 held out test epochs. Classes proportions were maintained and the same epoch assignments were applied to all channel configurations and classifiers. Partitioning was not patient-disjoint, so epochs from one participant could have been in more than one subset. This means that the results are interpreted as internal epoch-level performance and are not used to infer generalization to unseen patients. Table 3 states the information about distribution of mentioned five categories of IEDs across five anatomical regions in the dataset.

**Table 3:** Distribution of IEDs across five anatomical regions in the dataset.

| **IED Region** | **Count** | **Percent (%)** |
| --- | --- | --- |
| Temporal | 700 | 27.844% |
| Generalized | 573 | 22.792% |
| Frontal | 458 | 18.218% |
| Occipital | 417 | 16.587% |
| Centro-parietal | 366 | 14.558% |
| Total | 2514 | 100.000% |

### Addressing Class Imbalance with SMOTE Analysis

Synthetic Minority Over-sampling Technique (SMOTE) was applied only to the training subset to reduce imbalance among the five IED categories [21]. Validation and test epochs were not resampled. Applying SMOTE after the split prevented synthetic information from entering the validation or test sets.

For a minority-class feature vector *x_i_*, a synthetic vector was generated by interpolation between *x_i_* and a randomly selected nearest neighbour x_nn from the same class:

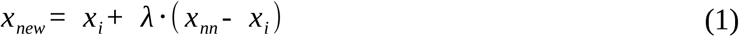

where:

- *x_i_* is the original minority-class sample;
- *x_nn_* is a randomly selected nearest neighbour from the same class;
- Lambda *λ* is sampled from a uniform distribution on [0,1]; and
- *x_new_* is the synthetic sample on the line segment joining *x_i_* and *x_nn_*.

### Machine Learning Classifier Evaluation

The classifiers evaluated are XGBoost, LightGBM, CatBoost, multinomial logistic regression, linear discriminant analysis (LDA), decision tree, extra trees, random forest and support vector machine (SVM). The random-forest analysis was available for the EEG-only and EEG with ECG analysis, while eight classifiers had results for all three channel configurations. The same procedure and partitions were utilized for each ablation. Hyperparameters were selected by using the validation set and the final results were reported on the held-out test set.

The performance was summarized in terms of overall accuracy, class-specific precision, recall and F1 score along with macro-averaged F1 score [22]. Macro-F1 was highlighted due to the fact that it gives equal importance to each spatial category irrespective of the class frequencies.

### Channel-Set Ablation Design

The channel-set ablation compared feature matrices derived from identical epochs. The EEG-only matrix contained 26 features from each of 19 scalp EEG channels. The EEG with ECG matrix added ECG RA and ECG LA. The full matrix contained all 29 recorded channels. Labels, split assignments, SMOTE based training, classifier families, and evaluation procedures were held constant. Consequently, the EEG-versus-EEG with ECG comparison estimates the numerical change associated with ECG-derived features. The EEG with ECG-versus-full comparison reflects the joint addition of four referential and four EMG channels and cannot isolate either group.

Performance differences between channel configurations are reported descriptively. Formal paired hypothesis tests and confidence intervals were not calculated because epoch-level prediction outputs required for paired error analysis were not retained for every configuration. Therefore, numerical differences should not be interpreted as statistically significant effects[23].

An additional exploratory EEG with EMG configuration, comprising the 19 scalp EEG channels and four bilateral deltoid EMG channels, was evaluated as a supplementary analysis. The same preprocessing, feature-extraction, data-partitioning, SMOTE based training, classifier-evaluation, and SHAP-analysis procedures were applied. Detailed results are provided in S1 File.

### SHapley Additive exPlanations (SHAP) Explainability Analysis

The contribution of each channel-feature variable to model output was described using SHAP values. Global importance was calculated by taking the average of absolute SHAP values over test epochs and classes, and then scaling it to percentages. These percentages represent a relative model dependence for a given feature set. They are not ‘effect sizes’, but rather ‘physiological effect sizes’, and as such should not be compared across configurations in a manner that assumes they measure the same causal quantity.

#### Shapley Value for a Feature *i*

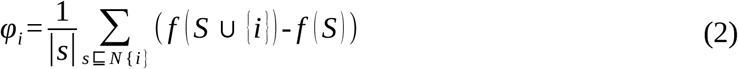

- For feature *i*, the Shapley value is the weighted average change in model output produced by adding that feature to all possible subsets of the remaining features.
- The calculation considers feature subsets that do not contain *i* and compares the prediction before and after *i* is added.
- The difference between these predictions is the marginal contribution of feature *i*.
- Weighted averaging across subsets yields the Shapley value.

#### Multi-Class SHAP (per sample and class)

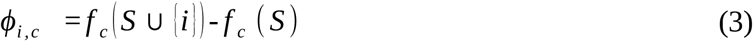

For multi-class classification, SHAP values are obtained for each sample, feature, and output class. A positive or negative SHAP value indicates whether the feature shifts the model output toward or away from a class. Class-specific SHAP values were converted to absolute magnitudes for global importance summaries. This formulation extends Shapley attribution to the five-class output.

#### SHAP Importance for Feature *i*

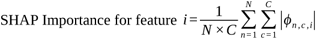

The overall significance of feature has been taken as interest here rather than a single sample or class. Thus, the absolute SHAP values are averaged across all *N* samples (data points) all *c* classes. Taking the absolute value makes sure both positive influence and negative influence are counted as importance. This gives a global importance score for each feature.

#### Normalization of SHAP Importance

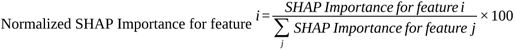

- The raw SHAP importance values might be arbitrary numbers.
- To make them comparable, we scale them so that:
- Now each feature’s importance is expressed as a percentage of total importance.

### Model Performance Metrics

Model performance was derived from the multi-class confusion matrix. For each class, true positives, false positives, and false negatives were defined using a one-versus-rest formulation. Overall accuracy measured the proportion of correctly classified epochs.

- True positives (TP): epochs of a class correctly assigned to that class.
- True negatives (TN): epochs outside a class correctly not assigned to that class.
- False positives (FP): epochs from other classes incorrectly assigned to the class.
- False negatives (FN): epochs of the class incorrectly assigned elsewhere. Precision, recall, and F1-score were calculated separately for each class.
- Accuracy was calculated as the number of correct predictions divided by the total number of test epochs.

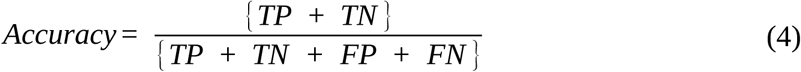

- Precision was calculated as TP/(TP+FP) and measures the proportion of predictions for a class that were correct.

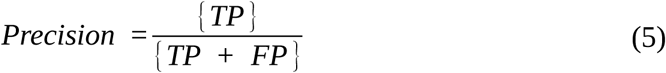

- Recall was calculated as TP/(TP+FN) and measures the proportion of epochs from a class that were correctly identified.

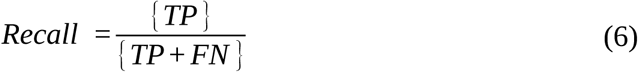

- F1-score was calculated as the harmonic mean of precision and recall. Macro-F1 was the unweighted mean of the five class-specific F1-scores.

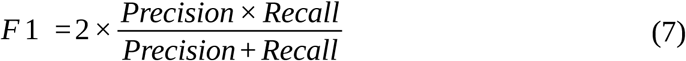

Together, these metrics summarize overall and class-balanced conditional classification performance.

## Results

Class-wise metrics were calculated for training, validation, and held-out test subsets. Model selection used the validation set, while principal conclusions are based on the untouched test set. Table 4 summarizes the original full 29-channel results. Tables 5 and 6 compare test accuracy and macro-F1 across channel configurations, and Table 7 reports CatBoost class-specific F1-scores. All between-configuration differences are descriptive.

**Table 4.** Full 29-channel test-set precision, recall, F1-score, and overall accuracy. For the full 29-channel configuration, CatBoost produced the highest test accuracy (94.44%) and macro-F1 (94.44%). Its class-specific F1-scores were 0.965 for generalized, 0.903 for frontal, 0.943 for temporal, 0.951 for occipital, and 0.960 for centro-parietal epochs. These values quantify conditional categorization among expert-confirmed IED epochs and are not measures of IED detection.

| **Metric** | **IED spatial category** | **XGB** | **LGB** | **CatB** | **LR** | **LDA** | **DT** | **ET** | **SVM** |
| --- | --- | --- | --- | --- | --- | --- | --- | --- | --- |
| Precision | Generalized | 0.960 | 0.941 | 0.982 | 0.800 | 0.959 | 0.767 | 0.926 | 0.708 |
|  | Frontal | 0.830 | 0.830 | 0.894 | 0.700 | 0.796 | 0.592 | 0.780 | 0.500 |
|  | Temporal | 0.880 | 0.892 | 0.943 | 0.870 | 0.857 | 0.848 | 0.873 | 0.780 |
|  | Occipital | 0.884 | 0.884 | 0.975 | 0.939 | 0.975 | 0.861 | 0.974 | 0.778 |
|  | Centro-parietal | 0.973 | 0.973 | 0.923 | 0.850 | 0.946 | 0.780 | 0.897 | 0.875 |
| Recall | Generalized | 0.828 | 0.828 | 0.948 | 0.828 | 0.810 | 0.793 | 0.862 | 0.793 |
|  | Frontal | 0.848 | 0.848 | 0.913 | 0.761 | 0.848 | 0.630 | 0.848 | 0.565 |
|  | Temporal | 0.943 | 0.943 | 0.943 | 0.857 | 0.943 | 0.800 | 0.886 | 0.557 |
|  | Occipital | 0.905 | 0.905 | 0.929 | 0.738 | 0.929 | 0.738 | 0.881 | 0.833 |
|  | Centro-parietal | 1.000 | 1.000 | 1.000 | 0.944 | 0.972 | 0.889 | 0.972 | 0.972 |
| F1 Score | Generalized | 0.889 | 0.881 | 0.965 | 0.814 | 0.879 | 0.780 | 0.893 | 0.748 |
|  | Frontal | 0.839 | 0.839 | 0.903 | 0.729 | 0.821 | 0.611 | 0.812 | 0.531 |
|  | Temporal | 0.910 | 0.917 | 0.943 | 0.863 | 0.898 | 0.824 | 0.879 | 0.650 |
|  | Occipital | 0.894 | 0.894 | 0.951 | 0.827 | 0.951 | 0.795 | 0.925 | 0.805 |
|  | Centro-parietal | 0.986 | 0.986 | 0.960 | 0.895 | 0.959 | 0.831 | 0.933 | 0.921 |
| Overall Accuracy |  | 0.9008 | 0.9008 | 0.9444 | 0.8254 | 0.8968 | 0.7698 | 0.8849 | 0.7183 |

**Table 5.** Test accuracy across channel configurations and descriptive differences (pp, percentage points).

| **Classifier** | **EEG only (%)** | **EEG with ECG (%)** | **Full 29-channel (%)** | **ECG gain vs EEG (pp)** | **Full gain vs EEG with ECG (pp)** |
| --- | --- | --- | --- | --- | --- |
| CatBoost | 86.90 | 93.25 | 94.44 | +6.35 | +1.19 |
| LightGBM | 85.71 | 88.10 | 90.08 | +2.39 | +1.98 |
| XGBoost | 84.92 | 88.89 | 90.08 | +3.97 | +1.19 |
| Logistic Regression | 76.98 | 79.37 | 82.54 | +2.39 | +3.17 |
| LDA | 88.89 | 90.87 | 89.68 | +1.98 | -1.19 |
| Decision Tree | 77.38 | 79.76 | 76.98 | +2.38 | -2.78 |
| Extra Trees | 82.54 | 86.51 | 88.49 | +3.97 | +1.98 |
| SVM | 57.94 | 63.49 | 71.83 | +5.55 | +8.34 |

**Table 6.** Test macro-F1 across channel configurations.

| **Classifier** | **EEG only** | **EEG with ECG** | **Full 29-channel** |
| --- | --- | --- | --- |
| CatBoost | 0.872 | 0.933 | 0.944 |
| LightGBM | 0.861 | 0.883 | 0.903 |
| XGBoost | 0.853 | 0.890 | 0.904 |
| Logistic Regression | 0.769 | 0.794 | 0.826 |
| LDA | 0.896 | 0.913 | 0.902 |
| Decision Tree | 0.775 | 0.797 | 0.768 |
| Extra Trees | 0.835 | 0.871 | 0.888 |
| SVM | 0.591 | 0.646 | 0.731 |

**Table 7.**
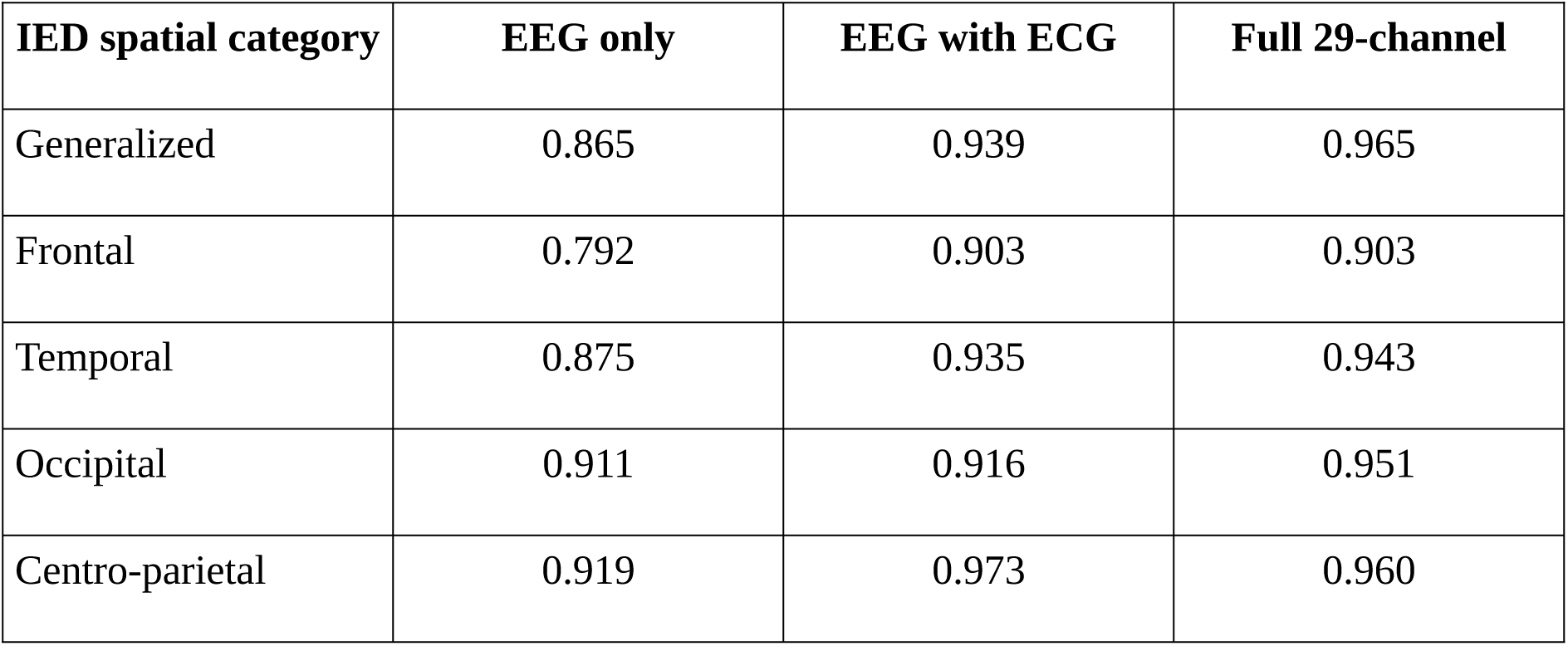
CatBoost class-specific test F1-scores across channel configurations. Relative to EEG-only CatBoost, EEG with ECG yielded higher F1-scores for generalized, frontal, temporal, occipital, and centro-parietal categories. The full configuration yielded the highest generalized, temporal, and occipital F1-scores; EEG with ECG yielded the highest centro-parietal F1-score; and frontal F1 was 0.903 in both ECG-containing configurations. These observations describe the fixed test split and were not subjected to formal inferential testing.

### Channel-Set Ablation Results

Adding ECG produced numerically higher held-out test accuracy for all eight classifiers evaluated across the three configurations. CatBoost increased from 86.90% on EEG channels only to 93.25% on EEG with ECG, a difference of 6.35 percentage points. The full 29-channel CatBoost result was 94.44%, a further numerical increase of 1.19 percentage points. The full configuration exceeded EEG with ECG for six classifiers, whereas LDA and decision tree performed better with EEG with ECG. Thus, the ECG addition was consistently associated with higher internal epoch-level accuracy, while the effect of the remaining channel groups was model-dependent.

Detailed validation- and test-set results for the EEG with EMG, EEG-only, and EEG with ECG configurations are provided in S1 File. These supplementary results include class-specific precision, recall, and F1-score, together with overall classification accuracy for the evaluated classifiers. Table 5 depicts the test accuracy across channel configurations and descriptive differences for EEG signals only, EEG with ECG signals and full 29 channel signals while Table 6 summarizes macro-F1 across channel configurations of test set. And finally, Table 7 summarizes CatBoost class-specific test F1-scores across different channel configurations

### EEG with ECG SHAP Analysis

Within the EEG with ECG CatBoost model as stated in Table 8, ECG RA and ECG LA received 15.79% and 15.12% of normalized global SHAP attribution, respectively. Together they accounted for 30.90% of normalized attribution in that feature set. Beta-band power was the leading feature family (18.76%), followed by peak-to-peak amplitude (9.72%), theta power (8.08%), alpha-beta ratio (7.98%), and alpha power (7.28%). Table 8 lists the largest individual channel-feature attributions.

**Table 8.**
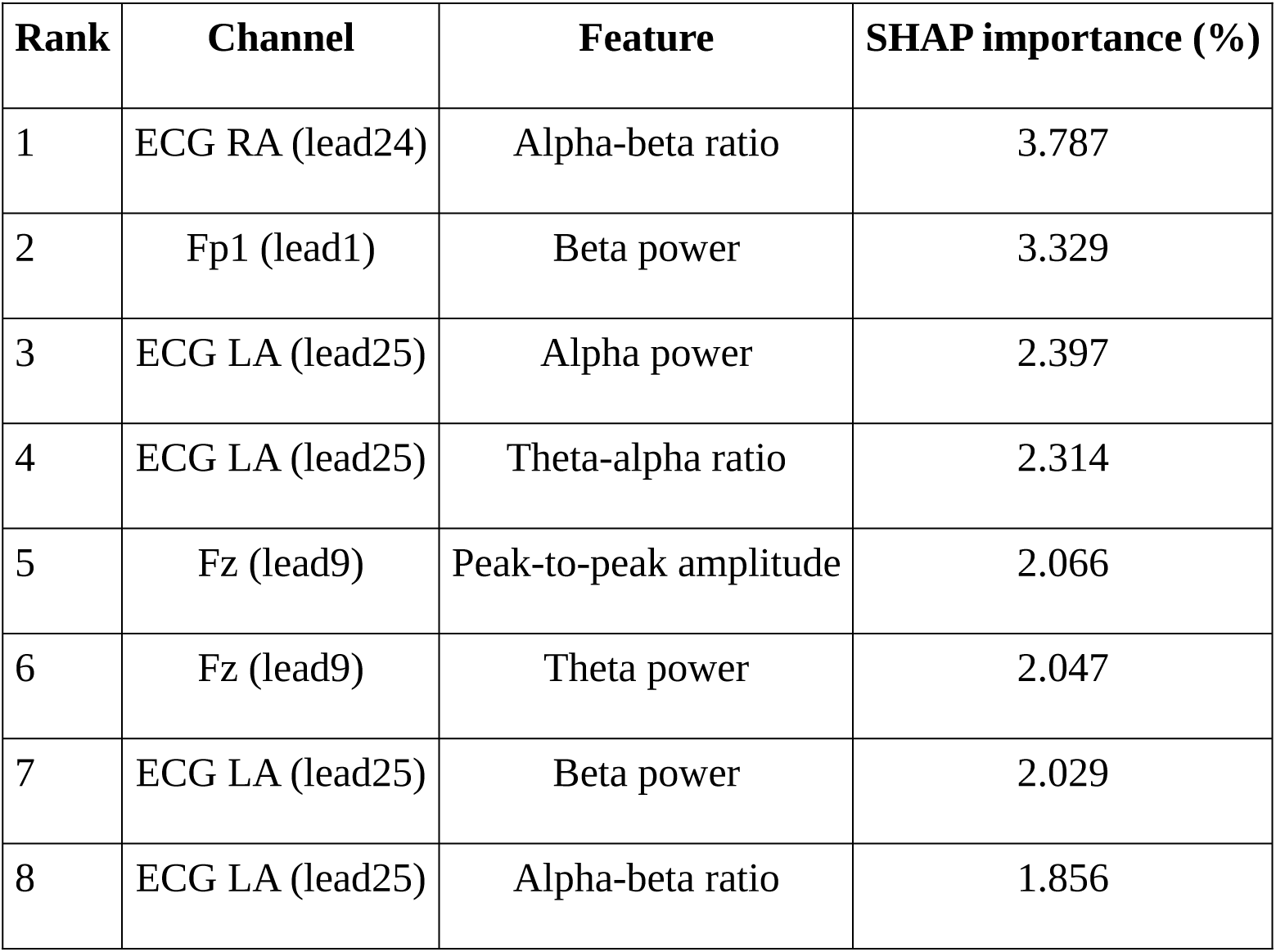
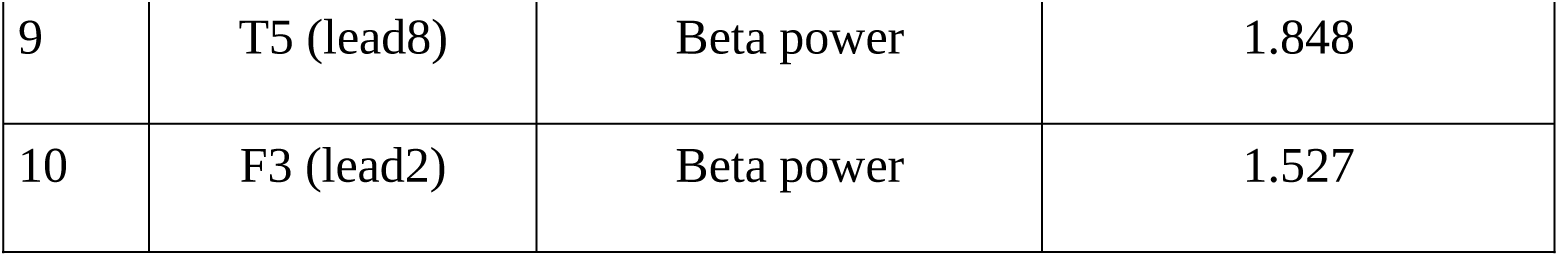
Largest CatBoost EEG with ECG channel-feature SHAP attributions on the test set. These values show that the fitted classifier relied strongly on ECG-derived variables within this dataset and target definition. They do not demonstrate that ECG contains a clinically meaningful interictal biomarker or autonomic localization signal. Patient identity, cardiac morphology, sleep-wake state, recording conditions, movement, contamination, and other dataset structure remain plausible alternative explanations.

| Rank | Channel | Feature | SHAP importance (%) |
| --- | --- | --- | --- |
| 1 | ECG RA (lead24) | Alpha-beta ratio | 3.787 |
| 2 | Fp1 (lead1) | Beta power | 3.329 |
| 3 | ECG LA (lead25) | Alpha power | 2.397 |
| 4 | ECG LA (lead25) | Theta-alpha ratio | 2.314 |
| 5 | Fz (lead9) | Peak-to-peak amplitude | 2.066 |
| 6 | Fz (lead9) | Theta power | 2.047 |
| 7 | ECG LA (lead25) | Beta power | 2.029 |
| 8 | ECG LA (lead25) | Alpha-beta ratio | 1.856 |
| 9 | T5 (lead8) | Beta power | 1.848 |
| 10 | F3 (lead2) | Beta power | 1.527 |

### SHAP Feature-Importance Analysis

Table 9 summarizes aggregated SHAP attribution for the full 29-channel models. The values identify variables used by the trained classifiers in the conditional five-class task; they do not show that a feature independently improved accuracy or represents a validated biomarker.

**Table 9:**
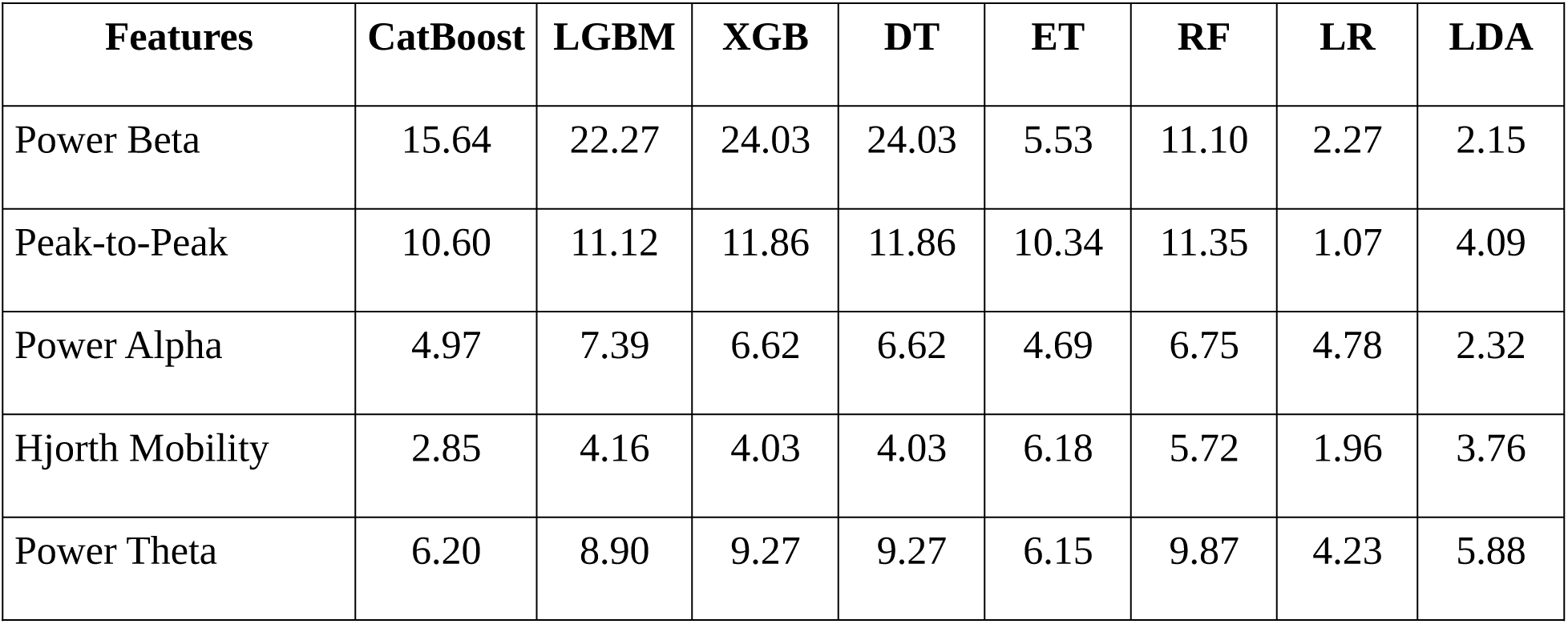
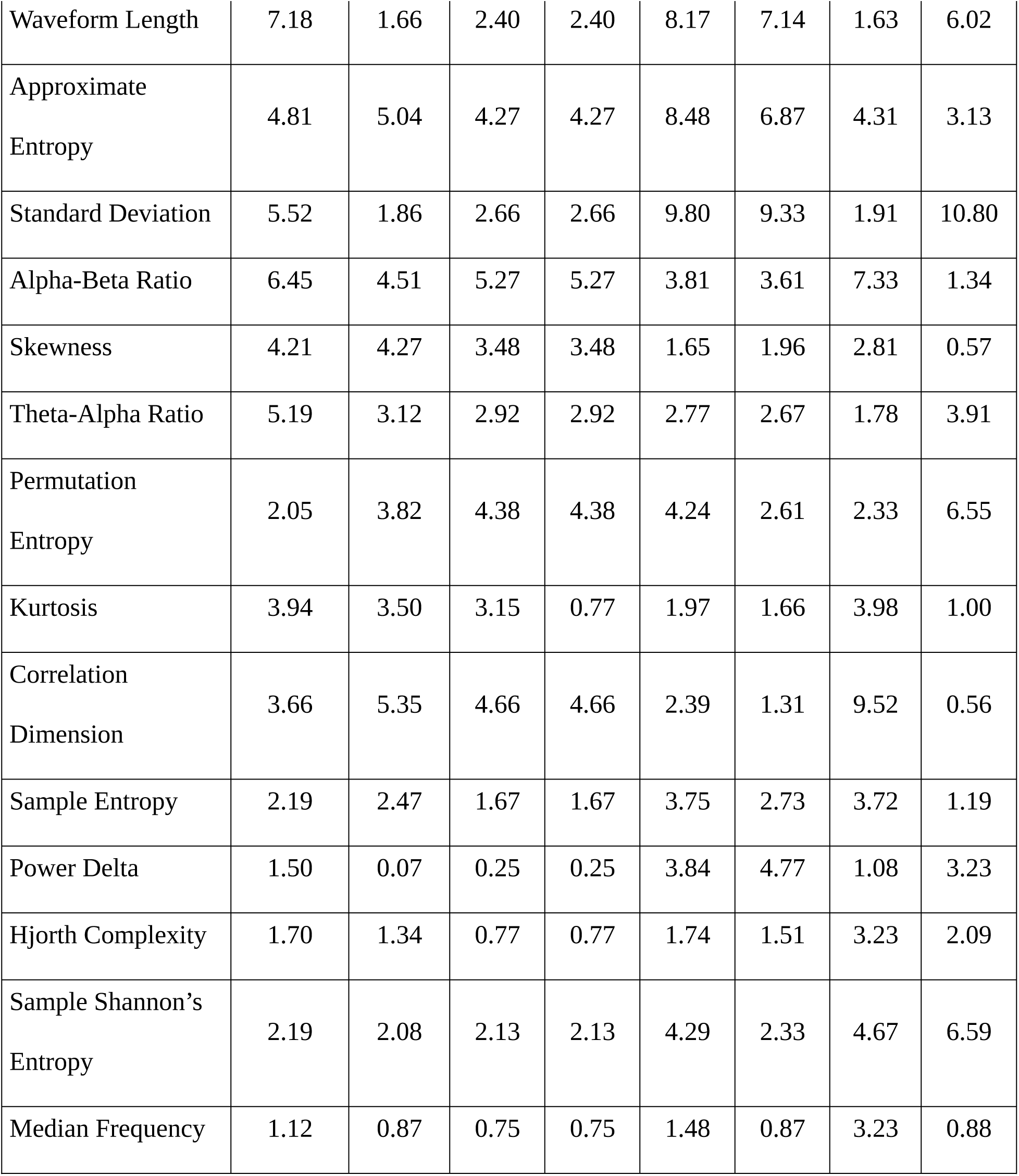
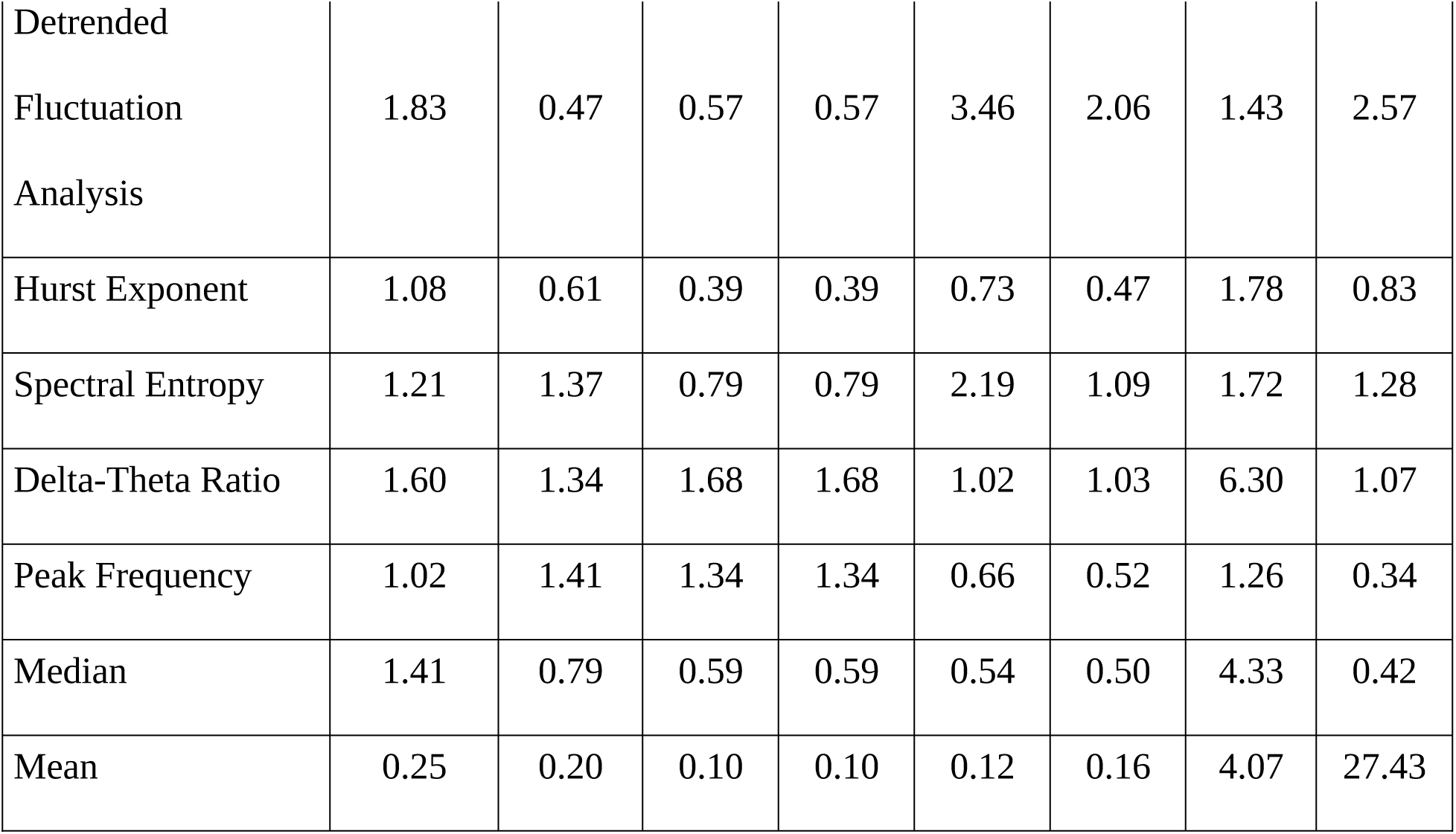
Aggregated feature SHAP importance across classifiers for the full 29-channel configuration.

| Features | CatBoost | LGBM | XGB | DT | ET | RF | LR | LDA |
| --- | --- | --- | --- | --- | --- | --- | --- | --- |
| Power Beta | 15.64 | 22.27 | 24.03 | 24.03 | 5.53 | 11.10 | 2.27 | 2.15 |
| Peak-to-Peak | 10.60 | 11.12 | 11.86 | 11.86 | 10.34 | 11.35 | 1.07 | 4.09 |
| Power Alpha | 4.97 | 7.39 | 6.62 | 6.62 | 4.69 | 6.75 | 4.78 | 2.32 |
| Hjorth Mobility | 2.85 | 4.16 | 4.03 | 4.03 | 6.18 | 5.72 | 1.96 | 3.76 |
| Power Theta | 6.20 | 8.90 | 9.27 | 9.27 | 6.15 | 9.87 | 4.23 | 5.88 |

| **Features** | **CatBoost** | **LGBM** | **XGB** | **DT** | **ET** | **RF** | **LR** | **LDA** |
| --- | --- | --- | --- | --- | --- | --- | --- | --- |
| Waveform Length | 7.18 | 1.66 | 2.40 | 2.40 | 8.17 | 7.14 | 1.63 | 6.02 |
| Approximate Entropy | 4.81 | 5.04 | 4.27 | 4.27 | 8.48 | 6.87 | 4.31 | 3.13 |
| Standard Deviation | 5.52 | 1.86 | 2.66 | 2.66 | 9.80 | 9.33 | 1.91 | 10.80 |
| Alpha-Beta Ratio | 6.45 | 4.51 | 5.27 | 5.27 | 3.81 | 3.61 | 7.33 | 1.34 |
| Skewness | 4.21 | 4.27 | 3.48 | 3.48 | 1.65 | 1.96 | 2.81 | 0.57 |
| Theta-Alpha Ratio | 5.19 | 3.12 | 2.92 | 2.92 | 2.77 | 2.67 | 1.78 | 3.91 |
| Permutation Entropy | 2.05 | 3.82 | 4.38 | 4.38 | 4.24 | 2.61 | 2.33 | 6.55 |
| Kurtosis | 3.94 | 3.50 | 3.15 | 0.77 | 1.97 | 1.66 | 3.98 | 1.00 |
| Correlation Dimension | 3.66 | 5.35 | 4.66 | 4.66 | 2.39 | 1.31 | 9.52 | 0.56 |
| Sample Entropy | 2.19 | 2.47 | 1.67 | 1.67 | 3.75 | 2.73 | 3.72 | 1.19 |
| Power Delta | 1.50 | 0.07 | 0.25 | 0.25 | 3.84 | 4.77 | 1.08 | 3.23 |
| Hjorth Complexity | 1.70 | 1.34 | 0.77 | 0.77 | 1.74 | 1.51 | 3.23 | 2.09 |
| Sample Shannon's Entropy | 2.19 | 2.08 | 2.13 | 2.13 | 4.29 | 2.33 | 4.67 | 6.59 |
| Median Frequency | 1.12 | 0.87 | 0.75 | 0.75 | 1.48 | 0.87 | 3.23 | 0.88 |
| Detrended<br>Fluctuation<br>Analysis | 1.83 | 0.47 | 0.57 | 0.57 | 3.46 | 2.06 | 1.43 | 2.57 |
| Hurst Exponent | 1.08 | 0.61 | 0.39 | 0.39 | 0.73 | 0.47 | 1.78 | 0.83 |
| Spectral Entropy | 1.21 | 1.37 | 0.79 | 0.79 | 2.19 | 1.09 | 1.72 | 1.28 |
| Delta-Theta Ratio | 1.60 | 1.34 | 1.68 | 1.68 | 1.02 | 1.03 | 6.30 | 1.07 |
| Peak Frequency | 1.02 | 1.41 | 1.34 | 1.34 | 0.66 | 0.52 | 1.26 | 0.34 |
| Median | 1.41 | 0.79 | 0.59 | 0.59 | 0.54 | 0.50 | 4.33 | 0.42 |
| Mean | 0.25 | 0.20 | 0.10 | 0.10 | 0.12 | 0.16 | 4.07 | 27.43 |

Table 10 reports channel-wise SHAP attribution for the full 29-channel configuration.

**Table 10:**
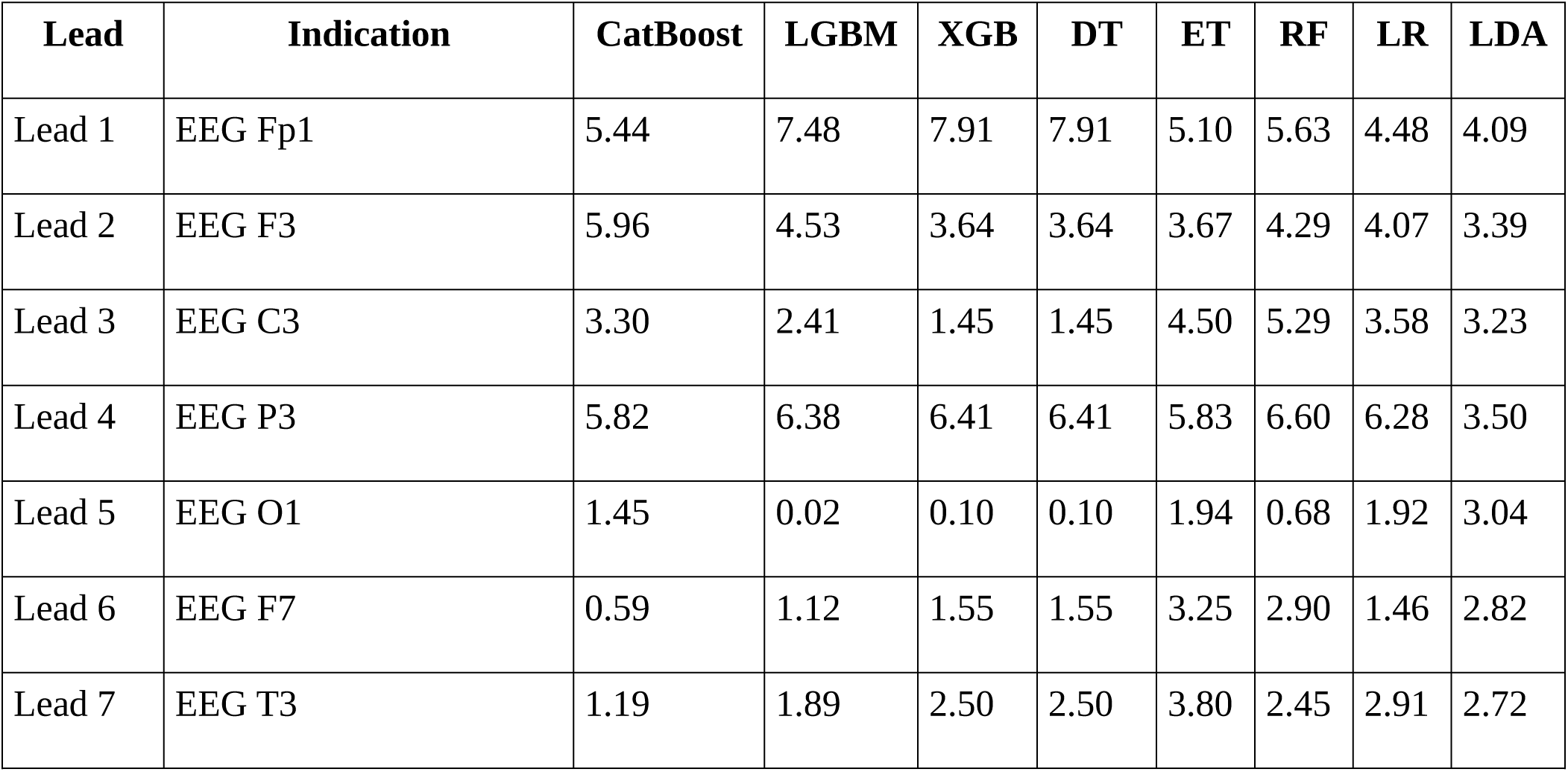
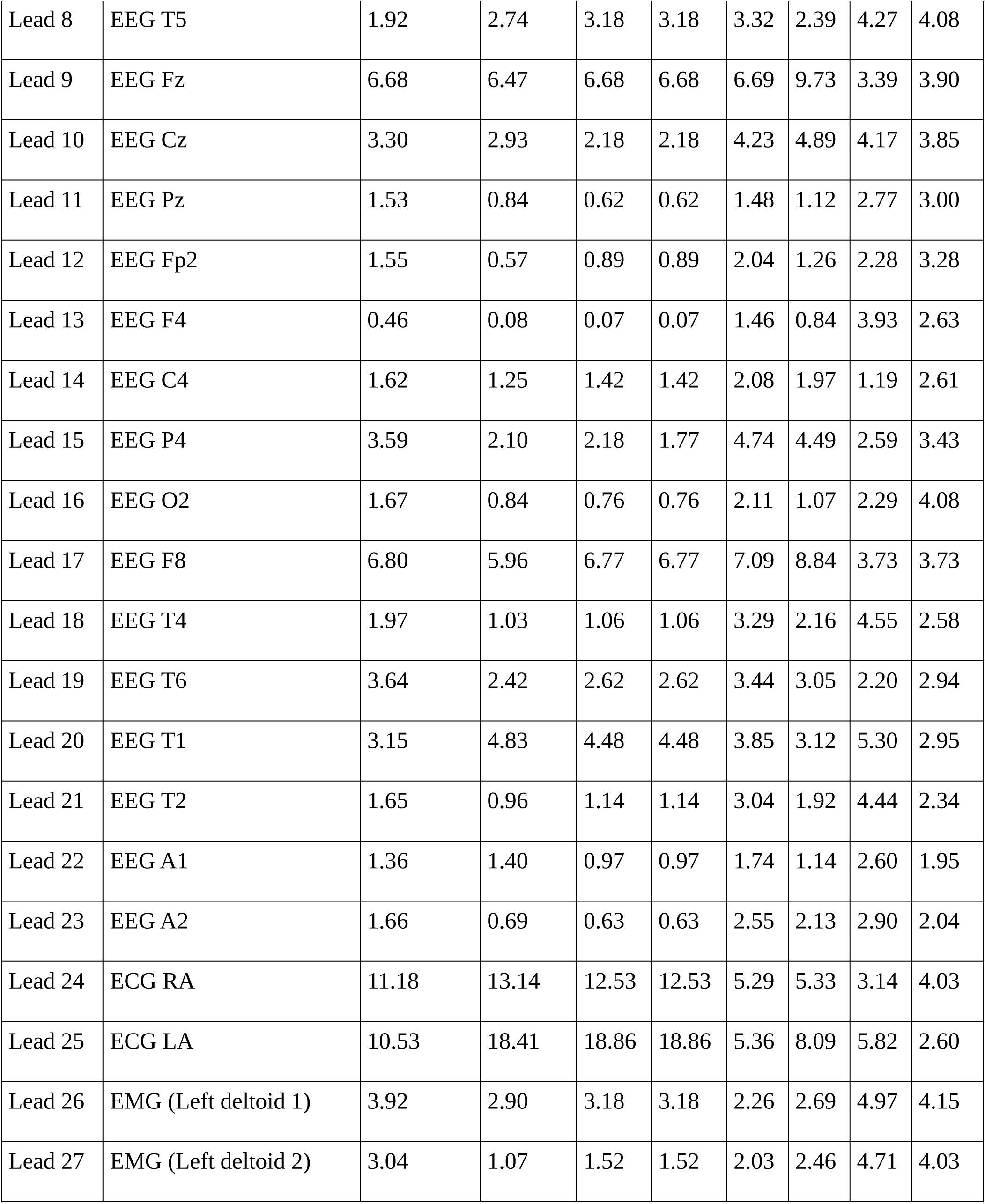
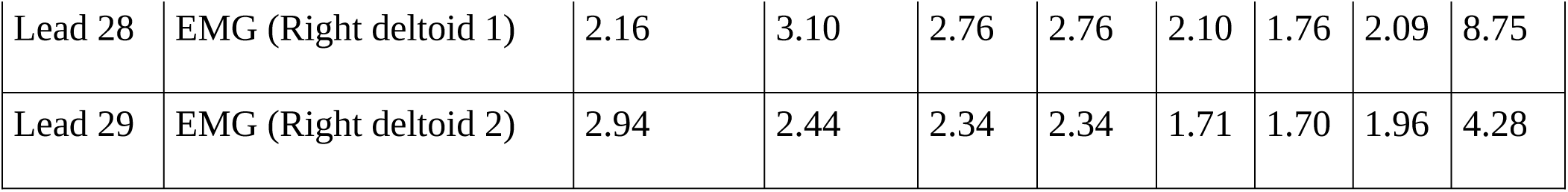
Channel-wise SHAP importance across classifiers for the full 29-channel configuration. CatBoost achieved the highest full-channel test accuracy. Table 11 lists its ten largest combined channel-feature attributions.

| Lead | Indication | CatBoost | LGBM | XGB | DT | ET | RF | LR | LDA |
| --- | --- | --- | --- | --- | --- | --- | --- | --- | --- |
| Lead 1 | EEG Fp1 | 5.44 | 7.48 | 7.91 | 7.91 | 5.10 | 5.63 | 4.48 | 4.09 |
| Lead 2 | EEG F3 | 5.96 | 4.53 | 3.64 | 3.64 | 3.67 | 4.29 | 4.07 | 3.39 |
| Lead 3 | EEG C3 | 3.30 | 2.41 | 1.45 | 1.45 | 4.50 | 5.29 | 3.58 | 3.23 |
| Lead 4 | EEG P3 | 5.82 | 6.38 | 6.41 | 6.41 | 5.83 | 6.60 | 6.28 | 3.50 |
| Lead 5 | EEG O1 | 1.45 | 0.02 | 0.10 | 0.10 | 1.94 | 0.68 | 1.92 | 3.04 |
| Lead 6 | EEG F7 | 0.59 | 1.12 | 1.55 | 1.55 | 3.25 | 2.90 | 1.46 | 2.82 |
| Lead 7 | EEG T3 | 1.19 | 1.89 | 2.50 | 2.50 | 3.80 | 2.45 | 2.91 | 2.72 |

| **Lead** | **Indication** | **CatBoost** | **LGBM** | **XGB** | **DT** | **ET** | **RF** | **LR** | **LDA** |
| --- | --- | --- | --- | --- | --- | --- | --- | --- | --- |
| Lead 8 | EEG T5 | 1.92 | 2.74 | 3.18 | 3.18 | 3.32 | 2.39 | 4.27 | 4.08 |
| Lead 9 | EEG Fz | 6.68 | 6.47 | 6.68 | 6.68 | 6.69 | 9.73 | 3.39 | 3.90 |
| Lead 10 | EEG Cz | 3.30 | 2.93 | 2.18 | 2.18 | 4.23 | 4.89 | 4.17 | 3.85 |
| Lead 11 | EEG Pz | 1.53 | 0.84 | 0.62 | 0.62 | 1.48 | 1.12 | 2.77 | 3.00 |
| Lead 12 | EEG Fp2 | 1.55 | 0.57 | 0.89 | 0.89 | 2.04 | 1.26 | 2.28 | 3.28 |
| Lead 13 | EEG F4 | 0.46 | 0.08 | 0.07 | 0.07 | 1.46 | 0.84 | 3.93 | 2.63 |
| Lead 14 | EEG C4 | 1.62 | 1.25 | 1.42 | 1.42 | 2.08 | 1.97 | 1.19 | 2.61 |
| Lead 15 | EEG P4 | 3.59 | 2.10 | 2.18 | 1.77 | 4.74 | 4.49 | 2.59 | 3.43 |
| Lead 16 | EEG O2 | 1.67 | 0.84 | 0.76 | 0.76 | 2.11 | 1.07 | 2.29 | 4.08 |
| Lead 17 | EEG F8 | 6.80 | 5.96 | 6.77 | 6.77 | 7.09 | 8.84 | 3.73 | 3.73 |
| Lead 18 | EEG T4 | 1.97 | 1.03 | 1.06 | 1.06 | 3.29 | 2.16 | 4.55 | 2.58 |
| Lead 19 | EEG T6 | 3.64 | 2.42 | 2.62 | 2.62 | 3.44 | 3.05 | 2.20 | 2.94 |
| Lead 20 | EEG T1 | 3.15 | 4.83 | 4.48 | 4.48 | 3.85 | 3.12 | 5.30 | 2.95 |
| Lead 21 | EEG T2 | 1.65 | 0.96 | 1.14 | 1.14 | 3.04 | 1.92 | 4.44 | 2.34 |
| Lead 22 | EEG A1 | 1.36 | 1.40 | 0.97 | 0.97 | 1.74 | 1.14 | 2.60 | 1.95 |
| Lead 23 | EEG A2 | 1.66 | 0.69 | 0.63 | 0.63 | 2.55 | 2.13 | 2.90 | 2.04 |
| Lead 24 | ECG RA | 11.18 | 13.14 | 12.53 | 12.53 | 5.29 | 5.33 | 3.14 | 4.03 |
| Lead 25 | ECG LA | 10.53 | 18.41 | 18.86 | 18.86 | 5.36 | 8.09 | 5.82 | 2.60 |
| Lead 26 | EMG (Left deltoid 1) | 3.92 | 2.90 | 3.18 | 3.18 | 2.26 | 2.69 | 4.97 | 4.15 |
| Lead 27 | EMG (Left deltoid 2) | 3.04 | 1.07 | 1.52 | 1.52 | 2.03 | 2.46 | 4.71 | 4.03 |
| Lead 28 | EMG (Right deltoid 1) | 2.16 | 3.10 | 2.76 | 2.76 | 2.10 | 1.76 | 2.09 | 8.75 |
| Lead 29 | EMG (Right deltoid 2) | 2.94 | 2.44 | 2.34 | 2.34 | 1.71 | 1.70 | 1.96 | 4.28 |

**Table 11:** Top combined channel-feature SHAP importance values for the full-channel CatBoost model. Across the full-channel models in Tables 9-11, spectral-power and waveform-morphology variables frequently received high attribution. Beta power and peak-to-peak amplitude were prominent in several tree-based classifiers. These results indicate that the models used these variables to separate the five expert-assigned labels; they do not establish that the variables detect IEDs or identify their physiological source.

| Rank | Feature | SHAP Importance (%) |
| --- | --- | --- |
| 1 | lead9_waveform_length | 2.68 |
| 2 | lead25_theta_alpha_ratio | 2.20 |
| 3 | lead19_power_beta | 2.11 |
| 4 | lead24_alpha_beta_ratio | 2.00 |
| 5 | lead17_ptp | 1.83 |
| 6 | lead1_power_beta | 1.74 |
| 7 | lead25_power_beta | 1.66 |
| 8 | lead25_power_alpha | 1.64 |
| 9 | lead17_power_beta | 1.52 |
| 10 | lead2_power_beta | 1.50 |

In the full configuration, ECG RA and ECG LA received high attribution in several classifiers, while scalp EEG channels including Fz, F8, and Fp1 also contributed. The ablation shows that adding ECG was accompanied by higher test accuracy across the eight comparable classifiers. Nevertheless, attribution alone does not establish a causal autonomic mechanism, and normalized SHAP percentages may change when the feature set changes.

In EEG-only CatBoost, beta power contributed 17.93% to normalized feature-family attribution and the most dominant channels were Fz (12.18%), F3 (10.74%), Fp1 (9.06%) and F8 (8.34%). In EEG with ECG CatBoost beta power was ranked 18.76%, while ECG RA and ECG LA were the two top ranked channels. These percentages are descriptive features of the fitted models.

Additional SHAP analyses are presented in S1 File, including aggregated feature-family importance, channel-wise importance, and the ten highest-ranked channel-feature attributions for each analysed classifier. These normalized SHAP values represent model-specific dependence within the corresponding feature set and should not be interpreted as physiological effect sizes, validated biomarkers, or evidence of causal mechanisms.

## Discussion

This study evaluated conditional five-class spatial categorization among expert-confirmed IED epochs and performed a staged channel-set ablation. Adding ECG was associated with numerically higher test accuracy for every directly comparable classifier, while the complete 29-channel input produced further gains for most, but not all, models. These findings apply only after IED presence has been established by expert annotation.

### Scope and Methodological Contribution

The retained task has a limited but defined methodological purpose. It tests whether an already identified IED epoch can be assigned to a standardized scalp-distribution category and whether auxiliary channel groups alter that categorization performance. This may be useful for benchmarking downstream annotation methods or organizing curated IED datasets, but it does not reproduce routine clinical EEG review, where IED presence is unknown. The manuscript therefore presents the work as an exploratory channel-ablation study rather than a clinically complete diagnostic system.

### Performance of the classifiers in the context of prior work

CatBoost test accuracy was 86.90% with EEG only, 93.25% with EEG with ECG, and 94.44% with the full 29-channel input. The numerical increase associated with adding ECG was larger than the subsequent increase associated with adding the remaining channels. However, the pattern depended on the classifier mainly LDA and decision tree performed better on EEG with ECG than with the full set. The results therefore support only a dataset-specific association between ECG inclusion and improved conditional classification; they do not establish a universal or causal benefit.

Table 12 places the present work alongside representative studies. Most prior reports address binary IED detection or use different spatial targets, datasets, and validation strategies. Their performance values are therefore not directly comparable with the present conditional five-class task.

**Table 12.**
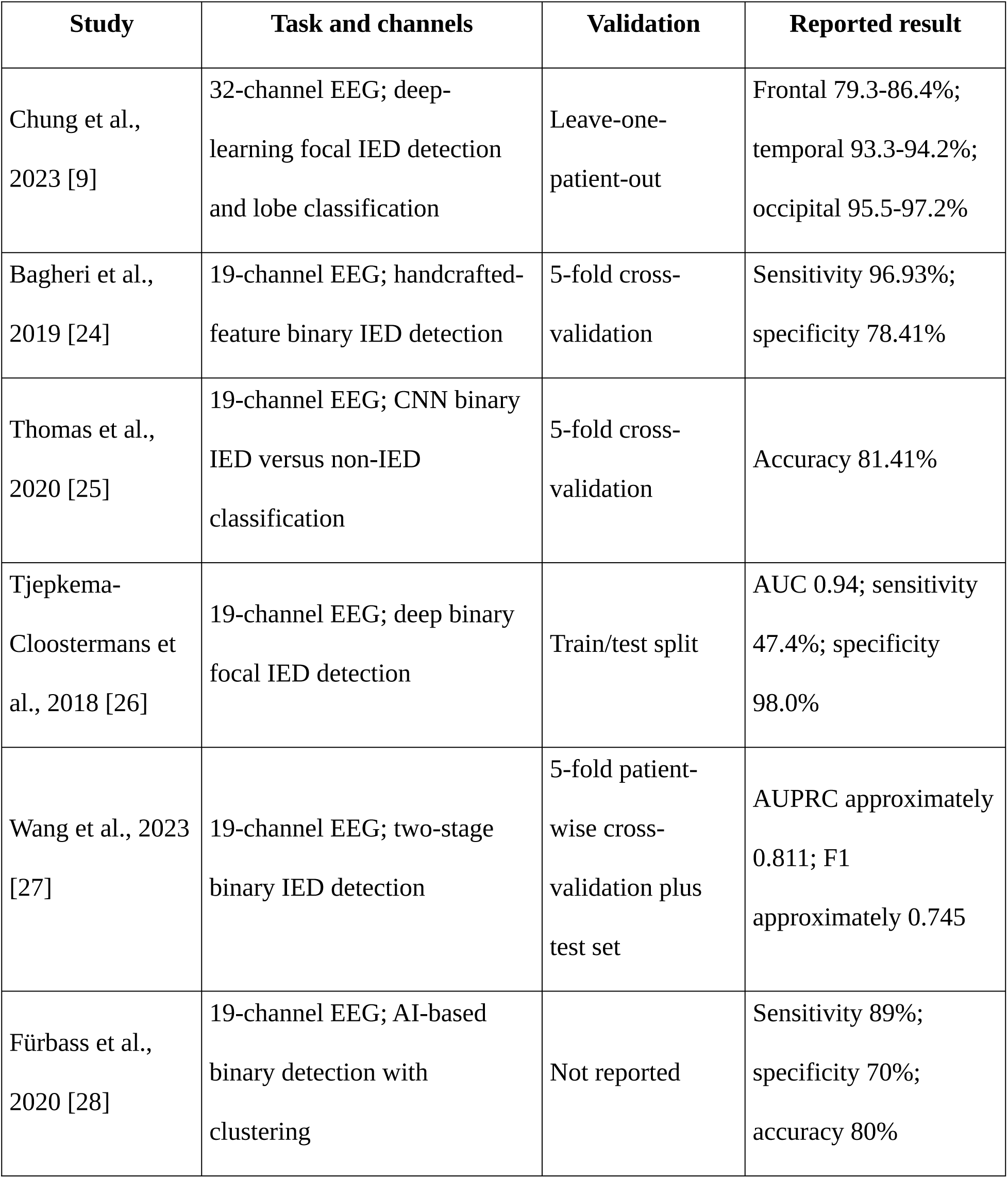
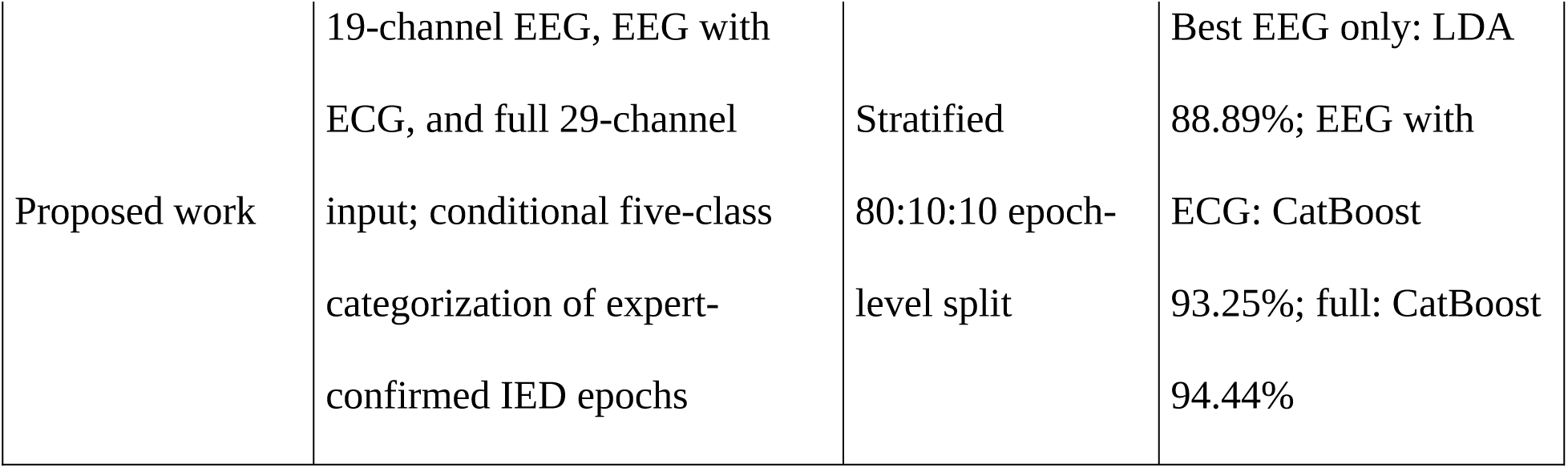
Comparison of the proposed framework with representative studies of IED detection and spatial classification. Previous studies indicate that performance may decline as epileptiform classification moves from binary decisions to finer spatial categories [6], [9]. The present results show that handcrafted-feature classifiers can separate five predefined labels within a curated collection of IED-positive epochs. This observation does not imply that the task is more clinically difficult than binary detection, because non-IED background, artifacts, and IED-like transients were excluded.

| Study | Task and channels | Validation | Reported result |
| --- | --- | --- | --- |
| Chung et al., 2023 [9] | 32-channel EEG; deep-learning focal IED detection and lobe classification | Leave-one-patient-out | Frontal 79.3-86.4%; temporal 93.3-94.2%; occipital 95.5-97.2% |
| Bagheri et al., 2019 [24] | 19-channel EEG; handcrafted-feature binary IED detection | 5-fold cross-validation | Sensitivity 96.93%; specificity 78.41% |
| Thomas et al., 2020 [25] | 19-channel EEG; CNN binary IED versus non-IED classification | 5-fold cross-validation | Accuracy 81.41% |
| Tjepkema-Cloostermans et al., 2018 [26] | 19-channel EEG; deep binary focal IED detection | Train/test split | AUC 0.94; sensitivity 47.4%; specificity 98.0% |
| Wang et al., 2023 [27] | 19-channel EEG; two-stage binary IED detection | 5-fold patient-wise cross-validation plus test set | AUPRC approximately 0.811; F1 approximately 0.745 |
| Fürbass et al., 2020 [28] | 19-channel EEG; AI-based binary detection with clustering | Not reported | Sensitivity 89%; specificity 70%; accuracy 80% |
| Proposed work | 19-channel EEG, EEG with ECG, and full 29-channel input; conditional five-class categorization of expert-confirmed IED epochs | Stratified 80:10:10 epoch-level split | Best EEG only: LDA 88.89%; EEG with ECG: CatBoost 93.25%; full: CatBoost 94.44% |

Classical and boosting-based methods remained competitive for the structured handcrafted features. CatBoost performed best with the ECG-containing configurations, whereas LDA performed best with EEG only. This configuration-dependent ranking argues against treating any model or channel set as universally superior.

Earlier feature-based studies showed that handcrafted EEG variables can support epileptiform analysis, often in binary detection settings [24]. The present contribution is narrower: it quantifies numerical changes from scalp EEG to EEG with ECG and then to the complete recorded channel set for conditional five-class categorization. It does not test whether these channels improve IED-versus-non-IED detection.

Cross-study comparisons are limited by differences in annotation, class definitions, preprocessing, sampling, validation, and channel composition. Moreover, the epoch-level split may place epochs from one participant in multiple subsets. The reported values should therefore be interpreted as internal held-out-epoch performance, not expected accuracy in unseen patients.

### Top Features and Leads (SHAP Analysis)

The ablation provides performance-based evidence that ECG-derived variables added information to the fitted models for this conditional task: all eight comparable classifiers showed numerically higher test accuracy with ECG than with scalp EEG alone. This result is stronger than a SHAP ranking alone, but it remains descriptive and dataset-specific. ECG was not evaluated for separating IED from non-IED activity, and the full-input comparison cannot identify an independent EMG effect because referential and EMG channels were added together.

SHAP provides a transparent account of variables used by a fitted model, not physiological validation. A frontal electrode may be important because it helps distinguish frontal-labelled epochs from other IED-labelled categories, which is partly expected from the target definition. Likewise, ECG ratios may contribute to class separation without being specific markers of IED presence or origin. The SHAP findings are therefore interpreted only as model attributions.

The prominence of beta power, waveform morphology, and selected EEG and ECG channels is consistent with the models exploiting spectral and amplitude differences among labelled epochs. This compatibility does not prove a physiological explanation. Normalized ECG attribution may also be influenced by feature-space composition. Patient-disjoint validation and controlled confound analyses are required before clinical or biological meaning can be assigned.

### Interpretation of ECG and EMG Contributions

High ECG attribution may reflect patient-specific cardiac morphology, sleep-wake state, medication, recording conditions, movement, cross-channel contamination, or other dataset structure; a genuine interictal neurocardiac association is only one possible explanation. The current analysis cannot distinguish among these alternatives. Seizure-related autonomic literature is therefore not treated as direct evidence for ECG-based spatial classification of interictal discharges.

The performance ablation shows that ECG inclusion was associated with improved conditional classification across the tested classifiers, but the mechanism remains unresolved. The full configuration does not isolate EMG because referential and EMG channels were introduced together. ECG SHAP prominence should therefore be regarded as hypothesis-generating rather than proof of an autonomic mechanism, a clinical biomarker, or spatial source information.

### Imbalanced Dataset Handling

SMOTE was only used on the training subset. Validation and test distributions were left unchanged. This reduced training imbalance but did not eliminate unequal class representation or guarantee equivalent performance across categories.

### Strengths of the Study

Strengths include expert-confirmed epoch labels, identical partitions across channel configurations, training-only oversampling, multiple classifier families, class-specific metrics, a staged channel ablation, and cautious SHAP interpretation. In particular, the EEG-versus-EEG with ECG comparison provides performance-based evidence about the incremental value of ECG within the retained conditional task rather than relying solely on feature importance.

### Limitations

Several limitations define the interpretation of this work. First, only expert-confirmed IED epochs were analyzed; consequently, the study does not evaluate IED detection, specificity, false-positive reduction, artifact rejection, or clinical screening. Second, the epoch-level split was not patient-disjoint. Participant overlap may allow models, particularly ECG-containing models, to exploit patient- or recording-specific information and may yield optimistic estimates relative to unseen-patient evaluation. Third, configuration differences are descriptive because formal paired tests and confidence intervals were not available. Fourth, although EEG with ECG isolates the addition of ECG, the full configuration jointly adds referential and EMG channels, so independent EMG and referential effects remain unknown. Fifth, spatial information is represented implicitly through channel-specific feature names rather than explicit electrode geometry or topographic maps. Finally, SHAP attributions are model-specific associations and do not establish physiological biomarkers or causal mechanisms.

### Future Directions

Future work should use patient-disjoint and external validation, preserve subject identifiers during feature extraction, and include representative non-IED background, artifacts, and IED-like transients. A clinically aligned extension could use a six-class formulation or a two-stage framework in which IED detection precedes spatial categorization. Separate ablations of referential, ECG, and EMG channels, together with explicit topographic or graph-based representations, would help determine whether auxiliary channels and electrode geometry provide reproducible value.

## Conclusion

This study presents an exploratory internal epoch-level analysis of conditional spatial classification among expert-confirmed IED epochs. Adding ECG to 19-channel scalp EEG was associated with numerically higher test accuracy for all eight directly comparable classifiers. CatBoost increased from 86.90% with EEG only to 93.25% with EEG with ECG and 94.44% with the complete 29-channel input. These results support a limited methodological conclusion that ECG-derived features contributed additional discriminatory information within this dataset and task. They do not establish automated IED detection, artifact rejection, independent EMG effects, physiological biomarkers, causal autonomic mechanisms, or generalization to unseen patients. Patient-disjoint validation and evaluation that includes non-IED epochs are required before clinical utility can be assessed.

## Supporting Information

Comprehensive supplementary performance and SHAP-attribution results are provided in S1 File. The file contains validation- and test-set analyses for the EEG with EMG, EEG-only, and EEG with ECG configurations, including class-specific performance metrics, aggregated feature-family importance, channel-wise importance, and classifier-specific channel-feature rankings.

## Data Availability Statement

No new data were collected. All source recordings and epoch labels are publicly available from Figshare at https://doi.org/10.6084/m9.figshare.28069568.v2.

## Code Availability Statement

The code used for preprocessing, feature extraction, model training, and SHAP analysis is available at https://github.com/Plabon826/EEG-Data-Processing-IED-Classification-SHAP-Feature-Importance-Analysis.

## Ethical Approval

This secondary analysis used a publicly available de-identified dataset and involved no new participant recruitment or data collection; additional institutional ethics approval was therefore not required.

## Consent to Participate

Not applicable because no participants were recruited for the present secondary analysis.

## Consent to Publish

Not applicable.

## Author Contributions (CRediT)

Conceptualization: Al Mukshit Plabon; Md. Faisal Mina

Methodology: Al Mukshit Plabon; Md. Faisal Mina; Torikul Islam

Software: Al Mukshit Plabon

Formal analysis: Al Mukshit Plabon

Investigation: Al Mukshit Plabon

Data curation: Al Mukshit Plabon; Abdul Mukit; Md. Neyamul; Omar Faruk Jehady; Fatima Tuz Zuba

Visualization: Al Mukshit Plabon; Abdul Mukit

Writing – original draft: Al Mukshit Plabon

Writing – review & editing: Al Mukshit Plabon; Md. Faisal Mina; Torikul Islam

Supervision: Md. Faisal Mina

Project administration: Md. Faisal Mina; Torikul Islam

## References

[1] K. Martin, C. F. Jackson, R. G. Levy, and P. N. Cooper, “Ketogenic diet and other dietary treatments for epilepsy,” Cochrane Database Syst. Rev., vol. 2, no. 2, Feb. 2016, doi: 10.1002/14651858.CD001903.PUB3.

[2] T. Islam et al., “Performance investigation of epilepsy detection from noisy EEG signals using base-2-meta stacking classifier,” Scientific Reports 2024 14:1, vol. 14, no. 1, pp. 10792-, May 2024, doi: 10.1038/s41598-024-61338-2.

[3] A. Davak, “A COMPREHENSIVE REVIEW ON EPILEPSY AND ITS DIAGNOSIS/TREATMENT USING EEG SIGNALS,” EPH - International Journal of Medical and Health Science, 2025, doi: 10.53555/EIJMHS.V11I1.267.

[4] T. Islam, M. Basak, R. Islam, and A. D. Roy, “Investigating population-specific epilepsy detection from noisy EEG signals using deep-learning models,” Heliyon, vol. 9, no. 12, p. e22208, Dec. 2023, doi: 10.1016/j.heliyon.2023.e22208.

[5] N. Lin et al., “An EEG dataset for interictal epileptiform discharge with spatial distribution information,” Sci. Data, vol. 12, no. 1, 2025, doi: 10.1038/S41597-025-04572-1.

[6] L. Zhang et al., “Automatic interictal epileptiform discharge (IED) detection based on convolutional neural network (CNN),” Front. Mol. Biosci., vol. 10, p. 1146606, Apr. 2023, doi: 10.3389/FMOLB.2023.1146606/BIBTEX.

[7] C. da Silva Lourenço, M. C. Tjepkema-Cloostermans, and M. J. A. M. van Putten, “Machine learning for detection of interictal epileptiform discharges,” Clinical Neurophysiology, vol. 132, no. 7, pp. 1433–1443, Jul. 2021, doi: 10.1016/J.CLINPH.2021.02.403.

[8] N. Lin et al., “vEpiNet: A multimodal interictal epileptiform discharge detection method based on video and electroencephalogram data,” Neural Networks, vol. 175, p. 106319, Jul. 2024, doi: 10.1016/J.NEUNET.2024.106319.

[9] Y. G. Chung et al., “Deep learning-based automated detection and multiclass classification of focal interictal epileptiform discharges in scalp electroencephalograms,” Sci. Rep., vol. 13, no. 1, Dec. 2023, doi: 10.1038/s41598-023-33906-5.

[10] S. Hooker, D. Erhan, P. J. Kindermans, and B. Kim, “A Benchmark for Interpretability Methods in Deep Neural Networks,” Adv. Neural Inf. Process. Syst., vol. 32, Jun. 2018, Accessed: Jul. 26, 2026. [Online]. Available: https://arxiv.org/pdf/1806.10758

[11] S. M. Lundberg and S. I. Lee, “A Unified Approach to Interpreting Model Predictions,” Adv. Neural Inf. Process. Syst., vol. 2017-December, pp. 4766–4775, May 2017, Accessed: Nov. 08, 2025. [Online]. Available: https://arxiv.org/pdf/1705.07874

[12] D. Janzing, L. Minorics, and P. Bloebaum, “Feature relevance quantification in explainable AI: A causal problem,” Jun. 03, 2020, PMLR. Accessed: Jul. 26, 2026. [Online]. Available: https://proceedings.mlr.press/v108/janzing20a.html

[13] W. Jin, X. Li, M. Fatehi, and G. Hamarneh, “Guidelines and evaluation of clinical explainable AI in medical image analysis,” Med. Image Anal., vol. 84, p. 102684, Feb. 2023, doi: 10.1016/J.MEDIA.2022.102684.

[14] S. Sanei and J. A. Chambers, “EEG signal processing and machine learning,” EEG Signal Processing and Machine Learning, pp. 1–752, Oct. 2021, doi: 10.1002/9781119386957.

[15] N. Bigdely-Shamlo, T. Mullen, C. Kothe, K. M. Su, and K. A. Robbins, “The PREP pipeline: Standardized preprocessing for large-scale EEG analysis,” Front. Neuroinform., vol. 9, no. JUNE, pp. 1–19, Jun. 2015, doi: 10.3389/FNINF.2015.00016/TEXT.

[16] A. Delorme and S. Makeig, “EEGLAB: An open source toolbox for analysis of single-trial EEG dynamics including independent component analysis,” J. Neurosci. Methods, vol. 134, no. 1, pp. 9–21, Mar. 2004, doi: 10.1016/j.jneumeth.2003.10.009.

[17] I. Stancin, M. Cifrek, and A. Jovic, “A review of eeg signal features and their application in driver drowsiness detection systems,” Sensors, vol. 21, no. 11, Jun. 2021, doi: 10.3390/S21113786.

[18] N. V. Chawla, K. W. Bowyer, L. O. Hall, and W. P. Kegelmeyer, “SMOTE: Synthetic Minority Over-sampling Technique,” Journal of Artificial Intelligence Research, vol. 16, pp. 321–357, Jun. 2002, doi: 10.1613/JAIR.953.

[19] R. Islam, S. Debnath, R. Raen, N. Islam, T. I. Palash, and R. Ali, “Epileptic Seizure Detection from EEG Signal Using ANN-LSTM Model,” Lecture Notes in Networks and Systems, vol. 675 LNNS, pp. 129–141, 2023, doi: 10.1007/978-981-99-1916-1_10.

[20] T. G. Dietterich, “Approximate Statistical Tests for Comparing Supervised Classification Learning Algorithms,” Neural Comput., vol. 10, no. 7, pp. 1895–1923, Oct. 1998, doi: 10.1162/089976698300017197.

[21] E. Bagheri, J. Jin, J. Dauwels, S. Cash, and M. B. Westover, “A fast machine learning approach to facilitate the detection of interictal epileptiform discharges in the scalp electroencephalogram,” J. Neurosci. Methods, vol. 326, Oct. 2019, doi: 10.1016/j.jneumeth.2019.108362.

[22] J. Thomas et al., “Automated Detection of Interictal Epileptiform Discharges from Scalp Electroencephalograms by Convolutional Neural Networks,” Int. J. Neural Syst., vol. 30, no. 11, Nov. 2020, doi: 10.1142/S0129065720500306,.

[23] M. C. Tjepkema-Cloostermans, R. C. V. de Carvalho, and M. J. A. M. van Putten, “Deep learning for detection of focal epileptiform discharges from scalp EEG recordings,” Clinical Neurophysiology, vol. 129, no. 10, pp. 2191–2196, Oct. 2018, doi: 10.1016/j.clinph.2018.06.024.

[24] X. Wang et al., “A Two-Stage Automatic System for Detection of Interictal Epileptiform Discharges from Scalp Electroencephalograms,” eNeuro, vol. 10, no. 11, Nov. 2023, doi: 10.1523/ENEURO.0111-23.2023.

[25] F. Fürbass, M. A. Kural, G. Gritsch, M. Hartmann, T. Kluge, and S. Beniczky, “An artificial intelligence-based EEG algorithm for detection of epileptiform EEG discharges: Validation against the diagnostic gold standard,” Clinical Neurophysiology, vol. 131, no. 6, pp. 1174–1179, Jun. 2020, doi: 10.1016/j.clinph.2020.02.032.

[26] Y. A. Saadoon, M. Khalil, and D. Battikh, “Machine and Deep Learning-Based Seizure Prediction: A Scoping Review on the Use of Temporal and Spectral Features,” Applied Sciences 2025 Vol. 15, Page 6279, vol. 15, no. 11, p. 6279, Jun. 2025, doi: 10.3390/APP15116279.

[27] R. Falach et al., “Annotated interictal discharges in intracranial EEG sleep data and related machine learning detection scheme,” Scientific Data, vol. 11, no. 1, Dec. 2024, doi: 10.1038/s41597-024-04187-y.

