## Supplementary material for "Conditional Spatial Classification of Expert-Confirmed Interictal Epileptiform Discharge Epochs: An EEG-ECG Ablation and SHAP Analysis": S1 file

### Supporting Information

Complete supplementary-table file for the manuscript "Conditional Spatial Classification of Expert-Confirmed Interictal Epileptiform Discharge Epochs: An EEG-ECG Ablation and SHAP Analysis." The captions and descriptions below are followed by S1 Table, S2 Table, and S3 Table in full.

#### **S1 Table.** EEG+EMG validation/test performance and SHAP importance results.

This supporting file contains validation- and test-set performance metrics, aggregated feature-family and channel-wise SHAP importance, and model-specific top-ten channel-feature SHAP rankings for the EEG+EMG configuration. Tables A-L are presented as editable Word tables.

##### Internal tables in S1 Table

**Table A.** Validation-set classification performance for the EEG+EMG configuration. Reports class-specific precision, recall, F1 score, and accuracy for the five listed categories across the evaluated classifiers, together with overall accuracy.

**Table B.** Test-set classification performance for the EEG+EMG configuration. Reports class-specific precision, recall, F1 score, and accuracy for the five listed categories across the evaluated classifiers, together with overall accuracy.

**Table C.** Aggregated feature importance for the EEG+EMG configuration. Reports normalized SHAP importance percentages aggregated by feature family across the evaluated classifiers.

**Table D.** Channel-wise importance for the EEG+EMG configuration. Reports normalized SHAP importance percentages aggregated by recording channel across the evaluated classifiers.

**Table E.** Top channel-feature SHAP attributions for CatBoost. Ranks the ten channel-feature variables with the largest normalized SHAP importance values for the CatBoost model.

**Table F.** Top channel-feature SHAP attributions for LightGBM. Ranks the ten channel-feature variables with the largest normalized SHAP importance values for the LightGBM model.

**Table G.** Top channel-feature SHAP attributions for XGBoost. Ranks the ten channel-feature variables with the largest normalized SHAP importance values for the XGBoost model.

**Table H.** Top channel-feature SHAP attributions for the decision tree. Ranks the ten channel-feature variables with the largest normalized SHAP importance values for the decision-tree model.

**Table I.** Top channel-feature SHAP attributions for extra trees. Ranks the ten channel-feature variables with the largest normalized SHAP importance values for the extra-trees model.

**Table J.** Top channel-feature SHAP attributions for random forest. Ranks the ten channel-feature variables with the largest normalized SHAP importance values for the random-forest model.

**Table K.** Top channel-feature SHAP attributions for logistic regression. Ranks the ten channel-feature variables with the largest normalized SHAP importance values for the logistic-regression model.

**Table L.** Top channel-feature SHAP attributions for linear discriminant analysis. Ranks the ten channel-feature variables with the largest normalized SHAP importance values for the linear-discriminant-analysis model.

#### **S2 Table.** EEG-only validation/test performance and SHAP importance results.

This supporting file contains validation- and test-set performance metrics, aggregated feature-family and channel-wise SHAP importance, and model-specific top-ten channel-feature SHAP rankings for the EEG-only configuration. Tables A-L are presented as editable Word tables.

##### Internal tables in S2 Table

**Table A.** Validation-set classification performance for the EEG-only configuration. Reports class-specific precision, recall, F1 score, and accuracy for the five listed categories across the evaluated classifiers, together with overall accuracy.

**Table B.** Test-set classification performance for the EEG-only configuration. Reports class-specific precision, recall, F1 score, and accuracy for the five listed categories across the evaluated classifiers, together with overall accuracy.

**Table C.** Aggregated feature importance for the EEG-only configuration. Reports normalized SHAP importance percentages aggregated by feature family across the evaluated classifiers.

**Table D.** Channel-wise importance for the EEG-only configuration. Reports normalized SHAP importance percentages aggregated by recording channel across the evaluated classifiers.

**Table E.** Top channel-feature SHAP attributions for CatBoost. Ranks the ten channel-feature variables with the largest normalized SHAP importance values for the CatBoost model.

**Table F.** Top channel-feature SHAP attributions for LightGBM. Ranks the ten channel-feature variables with the largest normalized SHAP importance values for the LightGBM model.

**Table G.** Top channel-feature SHAP attributions for XGBoost. Ranks the ten channel-feature variables with the largest normalized SHAP importance values for the XGBoost model.

**Table H.** Top channel-feature SHAP attributions for the decision tree. Ranks the ten channel-feature variables with the largest normalized SHAP importance values for the decision-tree model.

**Table I.** Top channel-feature SHAP attributions for extra trees. Ranks the ten channel-feature variables with the largest normalized SHAP importance values for the extra-trees model.

**Table J.** Top channel-feature SHAP attributions for random forest. Ranks the ten channel-feature variables with the largest normalized SHAP importance values for the random-forest model.

**Table K.** Top channel-feature SHAP attributions for logistic regression. Ranks the ten channel-feature variables with the largest normalized SHAP importance values for the logistic-regression model.

**Table L.** Top channel-feature SHAP attributions for linear discriminant analysis. Ranks the ten channel-feature variables with the largest normalized SHAP importance values for the linear-discriminant-analysis model.

#### **S3 Table.** EEG+ECG validation/test performance and SHAP importance results.

This supporting file contains validation- and test-set performance metrics, aggregated feature-family and channel-wise SHAP importance, and model-specific top-ten channel-feature SHAP rankings for the EEG+ECG configuration. Tables A-L are presented as editable Word tables.

##### Internal tables in S3 Table

**Table A.** Validation-set classification performance for the EEG+ECG configuration. Reports class-specific precision, recall, F1 score, and accuracy for the five listed categories across the evaluated classifiers, together with overall accuracy.

**Table B.** Test-set classification performance for the EEG+ECG configuration. Reports class-specific precision, recall, F1 score, and accuracy for the five listed categories across the evaluated classifiers, together with overall accuracy.

**Table C.** Aggregated feature importance for the EEG+ECG configuration. Reports normalized SHAP importance percentages aggregated by feature family across the evaluated classifiers.

**Table D.** Channel-wise importance for the EEG+ECG configuration. Reports normalized SHAP importance percentages aggregated by recording channel across the evaluated classifiers.

**Table E.** Top channel-feature SHAP attributions for CatBoost. Ranks the ten channel-feature variables with the largest normalized SHAP importance values for the CatBoost model.

**Table F.** Top channel-feature SHAP attributions for LightGBM. Ranks the ten channel-feature variables with the largest normalized SHAP importance values for the LightGBM model.

**Table G.** Top channel-feature SHAP attributions for XGBoost. Ranks the ten channel-feature variables with the largest normalized SHAP importance values for the XGBoost model.

**Table H.** Top channel-feature SHAP attributions for the decision tree. Ranks the ten channel-feature variables with the largest normalized SHAP importance values for the decision-tree model.

**Table I.** Top channel-feature SHAP attributions for extra trees. Ranks the ten channel-feature variables with the largest normalized SHAP importance values for the extra-trees model.

**Table J.** Top channel-feature SHAP attributions for random forest. Ranks the ten channel-feature variables with the largest normalized SHAP importance values for the random-forest model.

**Table K.** Top channel-feature SHAP attributions for logistic regression. Ranks the ten channel-feature variables with the largest normalized SHAP importance values for the logistic-regression model.

**Table L.** Top channel-feature SHAP attributions for linear discriminant analysis. Ranks the ten channel-feature variables with the largest normalized SHAP importance values for the linear-discriminant-analysis model.

### S1 Table

#### EEG+EMG validation/test performance and SHAP importance results

This supporting file contains validation- and test-set performance metrics, aggregated feature-family and channel-wise SHAP importance, and model-specific top-ten channel-feature SHAP rankings for the EEG+EMG configuration. Tables A-L are presented as editable Word tables.

##### Contents and table descriptions

| Internal table | Description |
|---|---|
| Table A | Validation-set classification performance for the EEG+EMG configuration. Reports class-specific precision, recall, F1 score, and accuracy for the five listed categories across the evaluated classifiers, together with overall accuracy. |
| Table B | Test-set classification performance for the EEG+EMG configuration. Reports class-specific precision, recall, F1 score, and accuracy for the five listed categories across the evaluated classifiers, together with overall accuracy. |
| Table C | Aggregated feature importance for the EEG+EMG configuration. Reports normalized SHAP importance percentages aggregated by feature family across the evaluated classifiers. |
| Table D | Channel-wise importance for the EEG+EMG configuration. Reports normalized SHAP importance percentages aggregated by recording channel across the evaluated classifiers. |
| Table E | Top channel-feature SHAP attributions for CatBoost. Ranks the ten channel-feature variables with the largest normalized SHAP importance values for the CatBoost model. |
| Table F | Top channel-feature SHAP attributions for LightGBM. Ranks the ten channel-feature variables with the largest normalized SHAP importance values for the LightGBM model. |
| Table G | Top channel-feature SHAP attributions for XGBoost. Ranks the ten channel-feature variables with the largest normalized SHAP importance values for the XGBoost model. |
| Table H | Top channel-feature SHAP attributions for the decision tree. Ranks the ten channel-feature variables with the largest normalized SHAP importance values for the decision-tree model. |
| Table I | Top channel-feature SHAP attributions for extra trees. Ranks the ten channel-feature variables with the largest normalized SHAP importance values for the extra-trees model. |
| Table J | Top channel-feature SHAP attributions for random forest. Ranks the ten channel-feature variables with the largest normalized SHAP importance values for the random-forest model. |
| Table K | Top channel-feature SHAP attributions for logistic regression. Ranks the ten channel-feature variables with the largest normalized SHAP importance values for the logistic-regression model. |
| Table L | Top channel-feature SHAP attributions for linear discriminant analysis. Ranks the ten channel-feature variables with the largest normalized SHAP importance values for the linear-discriminant-analysis model. |

**Formatting note:** All tables remain editable, cell-based objects. Numerical values and source table text were retained; only layout, captions, and descriptive legends were added.

**Table A.** Validation-set classification performance for the EEG+EMG configuration.

| Performance Metrics | Seizure Onset Zone | XGB | LGB | CatB | LR | LDA | DT | ET | RF | SVM |
|---|---|---|---|---|---|---|---|---|---|---|
| **Precision** | Generalized | 0.891 | 0.909 | 0.909 | 0.778 | 0.898 | 0.880 | 0.833 | 0.833 | 0.683 |
| | Frontal | 0.708 | 0.739 | 0.766 | 0.609 | 0.723 | 0.556 | 0.617 | 0.620 | 0.542 |
| | temporal | 0.846 | 0.859 | 0.883 | 0.787 | 0.835 | 0.889 | 0.863 | 0.878 | 0.860 |
| | occipital | 1.000 | 1.000 | 1.000 | 0.780 | 0.872 | 0.680 | 0.939 | 0.970 | 0.667 |
| | Centro-parietal | 1.000 | 1.000 | 0.974 | 0.971 | 1.000 | 0.912 | 0.841 | 0.925 | 0.694 |
| **Recall** | Generalized | 0.860 | 0.877 | 0.877 | 0.737 | 0.772 | 0.772 | 0.789 | 0.789 | 0.719 |
| | Frontal | 0.739 | 0.739 | 0.783 | 0.609 | 0.739 | 0.652 | 0.630 | 0.674 | 0.565 |
| | temporal | 0.943 | 0.957 | 0.971 | 0.843 | 0.943 | 0.800 | 0.900 | 0.929 | 0.529 |
| | occipital | 0.829 | 0.854 | 0.829 | 0.780 | 0.829 | 0.829 | 0.756 | 0.780 | 0.829 |
| | Centro-parietal | 0.973 | 1.000 | 1.000 | 0.919 | 1.000 | 0.838 | 1.000 | 1.000 | 0.919 |
| **F1 Score** | Generalized | 0.875 | 0.893 | 0.893 | 0.757 | 0.830 | 0.822 | 0.811 | 0.811 | 0.701 |
| | Frontal | 0.723 | 0.739 | 0.774 | 0.609 | 0.731 | 0.600 | 0.624 | 0.646 | 0.553 |
| | temporal | 0.892 | 0.905 | 0.925 | 0.814 | 0.886 | 0.842 | 0.881 | 0.903 | 0.655 |
| | occipital | 0.907 | 0.921 | 0.907 | 0.780 | 0.850 | 0.747 | 0.838 | 0.865 | 0.739 |
| | Centro-parietal | 0.986 | 1.000 | 0.987 | 0.944 | 1.000 | 0.873 | 0.914 | 0.961 | 0.791 |
| **Accuracy** | Generalized | 0.860 | 0.877 | 0.877 | 0.737 | 0.772 | 0.772 | 0.789 | 0.789 | 0.719 |
| | Frontal | 0.739 | 0.739 | 0.783 | 0.609 | 0.739 | 0.652 | 0.630 | 0.674 | 0.565 |
| | temporal | 0.943 | 0.957 | 0.971 | 0.843 | 0.943 | 0.800 | 0.900 | 0.929 | 0.529 |
| | occipital | 0.829 | 0.854 | 0.829 | 0.780 | 0.829 | 0.829 | 0.756 | 0.780 | 0.829 |
| | Centro-parietal | 0.973 | 1.000 | 1.000 | 0.919 | 1.000 | 0.838 | 1.000 | 1.000 | 0.919 |
| **Overall Accuracy** | | 0.8725 | 0.8884 | 0.8964 | 0.7769 | 0.8566 | 0.7769 | 0.8167 | 0.8367 | 0.6853 |

**Description:** Reports class-specific precision, recall, F1 score, and accuracy for the five listed categories across the evaluated classifiers, together with overall accuracy.
**Abbreviations:** XGB, XGBoost; LGB, LightGBM; CatB, CatBoost; LR, logistic regression; LDA, linear discriminant analysis; DT, decision tree; ET, extra trees; RF, random forest; SVM, support vector machine.

**Table B.** Test-set classification performance for the EEG+EMG configuration.

| Performance Metrics | Seizure Onset Zone | XGB | LGB | CatB | LR | LDA | DT | ET | RF | SVM |
|---|---|---|---|---|---|---|---|---|---|---|
| Precision | Generalized | 0.943 | 0.961 | 0.981 | 0.827 | 0.944 | 0.906 | 0.940 | 0.941 | 0.683 |
| | Frontal | 0.800 | 0.776 | 0.833 | 0.600 | 0.784 | 0.569 | 0.723 | 0.736 | 0.591 |
| | temporal | 0.882 | 0.883 | 0.893 | 0.787 | 0.914 | 0.881 | 0.789 | 0.861 | 0.717 |
| | occipital | 1.000 | 0.972 | 0.974 | 0.739 | 0.975 | 0.761 | 0.973 | 0.973 | 0.558 |
| | Centro-parietal | 0.947 | 0.923 | 0.923 | 0.744 | 0.946 | 0.857 | 0.833 | 0.897 | 0.875 |
| Recall | Generalized | 0.862 | 0.845 | 0.879 | 0.741 | 0.879 | 0.828 | 0.810 | 0.828 | 0.741 |
| | Frontal | 0.870 | 0.826 | 0.870 | 0.652 | 0.870 | 0.630 | 0.739 | 0.848 | 0.565 |
| | temporal | 0.957 | 0.971 | 0.957 | 0.686 | 0.914 | 0.843 | 0.857 | 0.886 | 0.543 |
| | occipital | 0.833 | 0.833 | 0.881 | 0.810 | 0.929 | 0.833 | 0.857 | 0.857 | 0.690 |
| | Centro-parietal | 1.000 | 1.000 | 1.000 | 0.889 | 0.972 | 0.833 | 0.972 | 0.972 | 0.972 |
| F1 Score | Generalized | 0.901 | 0.899 | 0.927 | 0.782 | 0.911 | 0.865 | 0.870 | 0.881 | 0.711 |
| | Frontal | 0.833 | 0.800 | 0.851 | 0.625 | 0.825 | 0.598 | 0.731 | 0.788 | 0.578 |
| | temporal | 0.918 | 0.925 | 0.924 | 0.733 | 0.914 | 0.861 | 0.822 | 0.873 | 0.618 |
| | occipital | 0.909 | 0.897 | 0.925 | 0.773 | 0.951 | 0.795 | 0.911 | 0.911 | 0.617 |
| | Centro-parietal | 0.973 | 0.960 | 0.960 | 0.810 | 0.959 | 0.845 | 0.897 | 0.933 | 0.921 |
| Accuracy | Generalized | 0.862 | 0.845 | 0.879 | 0.741 | 0.879 | 0.828 | 0.810 | 0.828 | 0.741 |
| | Frontal | 0.870 | 0.826 | 0.870 | 0.652 | 0.870 | 0.630 | 0.739 | 0.848 | 0.565 |
| | temporal | 0.957 | 0.971 | 0.957 | 0.686 | 0.914 | 0.843 | 0.857 | 0.886 | 0.543 |
| | occipital | 0.833 | 0.833 | 0.881 | 0.810 | 0.929 | 0.833 | 0.857 | 0.857 | 0.690 |
| | Centro-parietal | 1.000 | 1.000 | 1.000 | 0.889 | 0.972 | 0.833 | 0.972 | 0.972 | 0.972 |
| **Overall Accuracy** | | 0.9048 | 0.8968 | 0.9167 | 0.7421 | 0.9087 | 0.7976 | 0.8413 | 0.8730 | 0.6786 |

**Description:** Reports class-specific precision, recall, F1 score, and accuracy for the five listed categories across the evaluated classifiers, together with overall accuracy.
**Abbreviations:** XGB, XGBoost; LGB, LightGBM; CatB, CatBoost; LR, logistic regression; LDA, linear discriminant analysis; DT, decision tree; ET, extra trees; RF, random forest; SVM, support vector machine.

**Table C.** Aggregated feature importance for the EEG+EMG configuration.

| Features | CatBoost | LGBM | XGB | DT | ET | RF | LR | LDA |
|---|---|---|---|---|---|---|---|---|
| **Power Beta** | 16.99 | 23.78 | 24.14 | 24.14 | 5.49 | 11.82 | ~0 | 2.13 |
| **Peak-to-Peak** | 9.38 | 12.02 | 12.29 | 12.29 | 10.90 | 12.65 | ~0 | 4.24 |
| **Power Alpha** | 5.78 | 4.72 | 5.38 | 5.38 | 4.52 | 6.18 | ~0 | 1.54 |
| **Hjorth Mobility** | 3.08 | 3.02 | 2.82 | 2.82 | 5.34 | 5.66 | ~0 | 3.46 |
| **Power Theta** | 7.28 | 12.28 | 12.29 | 12.29 | 6.08 | 10.39 | ~0 | 5.05 |
| **Waveform Length** | 6.77 | 2.99 | 3.83 | 3.83 | 9.11 | 7.93 | ~0 | 4.88 |
| **Approximate Entropy** | 4.67 | 5.70 | 4.96 | 4.96 | 8.94 | 6.38 | ~0 | 3.40 |
| **Standard Deviation** | 4.11 | 2.04 | 1.76 | 1.76 | 9.90 | 9.58 | ~0 | 9.89 |
| **Alpha-Beta Ratio** | 2.36 | 1.74 | 1.61 | 1.61 | 3.64 | 3.07 | 0.98 | 1.33 |
| **Skewness** | 4.50 | 5.45 | 5.44 | 5.44 | 1.31 | 1.58 | 0.30 | 0.52 |
| **Theta-Alpha Ratio** | 2.17 | 1.11 | 1.69 | 1.69 | 2.88 | 2.66 | 2.39 | 4.95 |
| **Permutation Entropy** | 3.71 | 3.35 | 4.31 | 4.31 | 4.77 | 2.65 | ~0 | 6.17 |
| **Kurtosis** | 7.32 | 4.84 | 4.72 | 4.72 | 1.95 | 1.99 | 3.93 | 0.93 |
| **Correlation Dimension** | 4.58 | 6.83 | 6.22 | 6.22 | 2.65 | 1.37 | 0.01 | 0.57 |
| **Sample Entropy** | 1.82 | 1.43 | 1.13 | 1.13 | 4.31 | 2.57 | 35.20 | 1.20 |
| **Power Delta** | 2.55 | 0.82 | 0.69 | 0.69 | 3.75 | 4.60 | ~0 | 2.63 |
| **Hjorth Complexity** | 2.60 | 2.17 | 1.03 | 1.03 | 1.21 | 1.29 | 0.41 | 1.65 |
| **Sample Shannon's Entropy** | 2.81 | 1.76 | 1.50 | 1.50 | 4.43 | 2.00 | 46.67 | 6.36 |
| **Median Frequency** | 2.22 | 1.37 | 1.53 | 1.53 | 1.30 | 0.78 | 1.03 | 0.67 |
| **Detrended Fluctuation Analysis** | 1.42 | 0.16 | 0.29 | 0.29 | 2.89 | 1.87 | 0.01 | 2.65 |
| **Hurst Exponent** | 0.74 | 0.28 | 0.55 | 0.55 | 0.64 | 0.41 | ~0 | 0.79 |
| **Spectral Entropy** | 0.68 | 0.25 | 0.15 | 0.15 | 1.89 | 0.81 | 0.22 | 1.27 |
| **Delta-Theta Ratio** | 0.30 | 0.39 | 0.58 | 0.58 | 0.83 | 0.84 | 7.51 | 1.10 |
| **Peak Frequency** | 0.64 | 0.74 | 0.61 | 0.61 | 0.48 | 0.25 | 1.25 | 0.27 |
| **Median** | 1.16 | 0.59 | 0.27 | 0.27 | 0.52 | 0.40 | ~0 | 0.35 |
| **Mean** | 0.25 | 0.05 | 0.08 | 0.08 | 0.14 | 0.17 | ~0 | 31.87 |

**Description:** Reports normalized SHAP importance percentages aggregated by feature family across the evaluated classifiers. **Abbreviations:** SHAP, SHapley Additive exPlanations; LGBM, LightGBM; XGB, XGBoost; DT, decision tree; ET, extra trees; RF, random forest; LR, logistic regression; LDA, linear discriminant analysis.

**Table D.** Channel-wise importance for the EEG+EMG configuration.

| Lead | Indication | CatBoost | LGBM | XGB | DT | ET | RF | LR | LDA |
|---|---|---|---|---|---|---|---|---|---|
| **Lead 1** | EEG Fp1 | 7.47 | 9.93 | 10.29 | 10.29 | 5.28 | 7.25 | 5.00 | 4.77 |
| **Lead 2** | EEG F3 | 8.65 | 6.41 | 6.20 | 6.20 | 4.47 | 5.73 | 6.33 | 4.08 |
| **Lead 3** | EEG C3 | 3.47 | 1.80 | 2.13 | 2.13 | 6.07 | 5.90 | 5.72 | 4.20 |
| **Lead 4** | EEG P3 | 5.38 | 6.98 | 8.03 | 8.03 | 7.44 | 7.88 | 7.15 | 4.34 |
| **Lead 5** | EEG O1 | 0.81 | 0.31 | 0.29 | 0.29 | 2.58 | 1.13 | 2.74 | 3.48 |
| **Lead 6** | EEG F7 | 3.14 | 2.27 | 2.26 | 2.26 | 3.87 | 4.03 | 2.62 | 3.40 |
| **Lead 7** | EEG T3 | 2.15 | 3.18 | 3.63 | 3.63 | 4.34 | 2.97 | 3.99 | 3.12 |
| **Lead 8** | EEG T5 | 2.77 | 3.68 | 4.16 | 4.16 | 4.19 | 3.53 | 4.95 | 4.17 |
| **Lead 9** | EEG Fz | 10.66 | 11.16 | 11.58 | 11.58 | 8.56 | 11.16 | 5.99 | 4.98 |
| **Lead 10** | EEG Cz | 3.97 | 3.66 | 2.66 | 2.66 | 5.84 | 4.93 | 4.91 | 4.90 |
| **Lead 11** | EEG Pz | 1.14 | 1.95 | 1.68 | 1.68 | 1.94 | 1.42 | 3.82 | 3.17 |
| **Lead 12** | EEG Fp2 | 2.72 | 1.58 | 1.64 | 1.64 | 2.98 | 1.98 | 2.15 | 3.61 |
| **Lead 13** | EEG F4 | 1.59 | 0.53 | 0.52 | 0.52 | 1.95 | 1.19 | 4.68 | 2.98 |
| **Lead 14** | EEG C4 | 2.99 | 2.37 | 2.37 | 2.37 | 3.13 | 3.37 | 3.65 | 3.05 |
| **Lead 15** | EEG P4 | 3.48 | 3.00 | 3.01 | 3.01 | 6.09 | 6.80 | 3.34 | 3.92 |
| **Lead 16** | EEG O2 | 2.40 | 1.72 | 1.88 | 1.88 | 2.59 | 1.61 | 3.50 | 4.43 |
| **Lead 17** | EEG F8 | 6.19 | 8.09 | 8.41 | 8.41 | 8.73 | 9.38 | 4.97 | 4.23 |
| **Lead 18** | EEG T4 | 3.15 | 2.26 | 2.34 | 2.34 | 4.44 | 3.25 | 5.28 | 3.40 |
| **Lead 19** | EEG T6 | 4.21 | 5.80 | 5.95 | 5.95 | 4.48 | 3.42 | 2.68 | 3.96 |
| **Lead 26** | EMG (Left deltoid 1) | 5.89 | 7.13 | 7.02 | 7.02 | 3.11 | 4.31 | 4.89 | 5.42 |
| **Lead 27** | EMG (Left deltoid 2) | 6.61 | 4.97 | 3.77 | 3.77 | 3.04 | 4.03 | 6.10 | 5.01 |
| **Lead 28** | EMG (Right deltoid 1) | 4.18 | 5.47 | 4.87 | 4.87 | 2.45 | 2.26 | 2.75 | 10.44 |
| **Lead 29** | EMG (Right deltoid 2) | 6.89 | 5.62 | 5.20 | 5.20 | 2.31 | 2.36 | 2.67 | 4.81 |

**Description:** Reports normalized SHAP importance percentages aggregated by recording channel across the evaluated classifiers. **Abbreviations:** SHAP, SHapley Additive exPlanations; LGBM, LightGBM; XGB, XGBoost; DT, decision tree; ET, extra trees; RF, random forest; LR, logistic regression; LDA, linear discriminant analysis.

**Table E.** Top channel-feature SHAP attributions for CatBoost.

| Rank | Feature | SHAP Importance (%) |
|---|---|---|
| **1** | lead9_power_theta | 2.31 |
| **2** | lead2_power_beta | 2.21 |
| **3** | lead19_power_beta | 2.16 |
| **4** | lead1_power_beta | 1.94 |
| **5** | lead9_waveform_length | 1.93 |
| **6** | lead17_power_beta | 1.68 |
| **7** | lead9_ptp | 1.66 |
| **8** | lead4_kurtosis | 1.35 |
| **9** | lead1_power_alpha | 1.27 |
| **10** | lead9_power_alpha | 1.23 |

**Description:** Ranks the ten channel-feature variables with the largest normalized SHAP importance values for the CatBoost model. **Abbreviations:** SHAP, SHapley Additive exPlanations.

**Table F.** Top channel-feature SHAP attributions for LightGBM.

| Rank | Feature | SHAP Importance (%) |
|---|---|---|
| **1** | lead9_power_theta | 4.72 |
| **2** | lead1_power_beta | 3.99 |
| **3** | lead26_power_beta | 2.80 |
| **4** | lead9_power_alpha | 2.63 |
| **5** | lead2_power_beta | 2.59 |
| **6** | lead19_power_beta | 2.06 |
| **7** | lead17_approx_entropy | 1.90 |
| **8** | lead4_power_theta | 1.88 |
| **9** | lead8_power_beta | 1.81 |
| **10** | lead17_power_beta | 1.79 |

**Description:** Ranks the ten channel-feature variables with the largest normalized SHAP importance values for the LightGBM model. **Abbreviations:** SHAP, SHapley Additive exPlanations.

**Table G.** Top channel-feature SHAP attributions for XGBoost.

| Rank | Feature | SHAP Importance (%) |
|---|---|---|
| **1** | lead9_power_theta | 4.99 |
| **2** | lead1_power_beta | 4.80 |
| **3** | lead9_power_alpha | 2.79 |
| **4** | lead19_power_beta | 2.37 |
| **5** | lead26_power_beta | 2.21 |
| **6** | lead2_power_beta | 2.17 |
| **7** | lead17_approx_entropy | 1.92 |
| **8** | lead4_power_theta | 1.89 |
| **9** | lead8_power_beta | 1.87 |
| **10** | lead17_ptp | 1.86 |

**Description:** Ranks the ten channel-feature variables with the largest normalized SHAP importance values for the XGBoost model. **Abbreviations:** SHAP, SHapley Additive exPlanations.

**Table H.** Top channel-feature SHAP attributions for the decision tree.

| Rank | Feature | SHAP Importance (%) |
|---|---|---|
| **1** | lead9_power_theta | 4.99 |
| **2** | lead1_power_beta | 4.80 |
| **3** | lead9_power_alpha | 2.79 |
| **4** | lead19_power_beta | 2.37 |
| **5** | lead26_power_beta | 2.21 |
| **6** | lead2_power_beta | 2.17 |
| **7** | lead17_approx_entropy | 1.92 |
| **8** | lead4_power_theta | 1.89 |
| **9** | lead8_power_beta | 1.87 |
| **10** | lead17_ptp | 1.86 |

**Description:** Ranks the ten channel-feature variables with the largest normalized SHAP importance values for the decision-tree model. **Abbreviations:** SHAP, SHapley Additive exPlanations.

**Table I.** Top channel-feature SHAP attributions for extra trees.

| Rank | Feature | SHAP Importance (%) |
|---|---|---|
| **1** | lead9_ptp | 1.17 |
| **2** | lead9_waveform_length | 1.14 |
| **3** | lead17_std | 1.09 |
| **4** | lead9_std | 0.99 |
| **5** | lead15_std | 0.98 |
| **6** | lead3_ptp | 0.96 |
| **7** | lead15_power_theta | 0.94 |
| **8** | lead17_ptp | 0.91 |
| **9** | lead10_waveform_length | 0.86 |
| **10** | lead4_std | 0.86 |

**Description:** Ranks the ten channel-feature variables with the largest normalized SHAP importance values for the extra-trees model. **Abbreviations:** SHAP, SHapley Additive exPlanations.

**Table J.** Top channel-feature SHAP attributions for random forest.

| Rank | Feature | SHAP Importance (%) |
|---|---|---|
| **1** | lead9_power_theta | 2.34 |
| **2** | lead9_ptp | 1.88 |
| **3** | lead15_power_theta | 1.83 |
| **4** | lead9_std | 1.52 |
| **5** | lead15_ptp | 1.50 |
| **6** | lead1_power_beta | 1.47 |
| **7** | lead9_power_alpha | 1.38 |
| **8** | lead2_power_beta | 1.35 |
| **9** | lead17_std | 1.32 |
| **10** | lead17_approx_entropy | 1.24 |

**Description:** Ranks the ten channel-feature variables with the largest normalized SHAP importance values for the random-forest model. **Abbreviations:** SHAP, SHapley Additive exPlanations.

**Table K.** Top channel-feature SHAP attributions for logistic regression.

| Rank | Feature | SHAP Importance (%) |
|---|---|---|
| **1** | lead4_ssc | 3.96 |
| **2** | lead2_ssc | 3.79 |
| **3** | lead3_ssc | 3.42 |
| **4** | lead9_ssc | 2.89 |
| **5** | lead27_ssc | 2.81 |
| **6** | lead8_sample_entropy | 2.68 |
| **7** | lead18_ssc | 2.48 |
| **8** | lead1_ssc | 2.30 |
| **9** | lead17_ssc | 2.28 |
| **10** | lead4_sample_entropy | 2.17 |

**Description:** Ranks the ten channel-feature variables with the largest normalized SHAP importance values for the logistic-regression model. **Abbreviations:** SHAP, SHapley Additive exPlanations.

**Table L.** Top channel-feature SHAP attributions for linear discriminant analysis.

| Rank | Feature | SHAP Importance (%) |
|---|---|---|
| 1 | lead28_theta_alpha_ratio | 3.52 |
| 2 | lead28_power_theta | 3.44 |
| 3 | lead2_mean | 2.44 |
| 4 | lead1_mean | 2.44 |
| 5 | lead10_mean | 2.08 |
| 6 | lead17_mean | 2.01 |
| 7 | lead9_mean | 2.01 |
| 8 | lead4_mean | 1.89 |
| 9 | lead16_mean | 1.82 |
| 10 | lead3_mean | 1.72 |

**Description:** Ranks the ten channel-feature variables with the largest normalized SHAP importance values for the linear-discriminant-analysis model. **Abbreviations:** SHAP, SHapley Additive exPlanations.

### S2 Table

#### EEG-only validation/test performance and SHAP importance results

This supporting file contains validation- and test-set performance metrics, aggregated feature-family and channel-wise SHAP importance, and model-specific top-ten channel-feature SHAP rankings for the EEG-only configuration. Tables A-L are presented as editable Word tables.

##### Contents and table descriptions

| Internal table | Description |
|---|---|
| Table A | Validation-set classification performance for the EEG-only configuration. Reports class-specific precision, recall, F1 score, and accuracy for the five listed categories across the evaluated classifiers, together with overall accuracy. |
| Table B | Test-set classification performance for the EEG-only configuration. Reports class-specific precision, recall, F1 score, and accuracy for the five listed categories across the evaluated classifiers, together with overall accuracy. |
| Table C | Aggregated feature importance for the EEG-only configuration. Reports normalized SHAP importance percentages aggregated by feature family across the evaluated classifiers. |
| Table D | Channel-wise importance for the EEG-only configuration. Reports normalized SHAP importance percentages aggregated by recording channel across the evaluated classifiers. |
| Table E | Top channel-feature SHAP attributions for CatBoost. Ranks the ten channel-feature variables with the largest normalized SHAP importance values for the CatBoost model. |
| Table F | Top channel-feature SHAP attributions for LightGBM. Ranks the ten channel-feature variables with the largest normalized SHAP importance values for the LightGBM model. |
| Table G | Top channel-feature SHAP attributions for XGBoost. Ranks the ten channel-feature variables with the largest normalized SHAP importance values for the XGBoost model. |
| Table H | Top channel-feature SHAP attributions for the decision tree. Ranks the ten channel-feature variables with the largest normalized SHAP importance values for the decision-tree model. |
| Table I | Top channel-feature SHAP attributions for extra trees. Ranks the ten channel-feature variables with the largest normalized SHAP importance values for the extra-trees model. |
| Table J | Top channel-feature SHAP attributions for random forest. Ranks the ten channel-feature variables with the largest normalized SHAP importance values for the random-forest model. |
| Table K | Top channel-feature SHAP attributions for logistic regression. Ranks the ten channel-feature variables with the largest normalized SHAP importance values for the logistic-regression model. |
| Table L | Top channel-feature SHAP attributions for linear discriminant analysis. Ranks the ten channel-feature variables with the largest normalized SHAP importance values for the linear-discriminant-analysis model. |

**Formatting note:** All tables remain editable, cell-based objects. Numerical values and source table text were retained; only layout, captions, and descriptive legends were added.

**Table A.** Validation-set classification performance for the EEG-only configuration.

| Performance Metrics | Seizure Onset Zone | XGB | LGB | CatB | LR | LDA | DT | ET | RF | SVM |
|---|---|---|---|---|---|---|---|---|---|---|
| Precision | Generalized | 0.891 | 0.909 | 0.909 | 0.778 | 0.898 | 0.880 | 0.833 | 0.833 | 0.683 |
| | Frontal | 0.708 | 0.739 | 0.766 | 0.609 | 0.723 | 0.556 | 0.617 | 0.620 | 0.542 |
| | Temporal | 0.846 | 0.859 | 0.883 | 0.787 | 0.835 | 0.889 | 0.863 | 0.878 | 0.860 |
| | Occipital | 1.000 | 1.000 | 1.000 | 0.780 | 0.872 | 0.680 | 0.939 | 0.970 | 0.667 |
| | Centro-parietal | 1.000 | 1.000 | 0.974 | 0.971 | 1.000 | 0.912 | 0.841 | 0.925 | 0.694 |
| Recall | Generalized | 0.860 | 0.877 | 0.877 | 0.737 | 0.772 | 0.772 | 0.789 | 0.789 | 0.719 |
| | Frontal | 0.739 | 0.739 | 0.783 | 0.609 | 0.739 | 0.652 | 0.630 | 0.674 | 0.565 |
| | Temporal | 0.943 | 0.957 | 0.971 | 0.843 | 0.943 | 0.800 | 0.900 | 0.929 | 0.529 |
| | Occipital | 0.829 | 0.854 | 0.829 | 0.780 | 0.829 | 0.829 | 0.756 | 0.780 | 0.829 |
| | Centro-parietal | 0.973 | 1.000 | 1.000 | 0.919 | 1.000 | 0.838 | 1.000 | 1.000 | 0.919 |
| F1 Score | Generalized | 0.875 | 0.893 | 0.893 | 0.757 | 0.830 | 0.822 | 0.811 | 0.811 | 0.701 |
| | Frontal | 0.723 | 0.739 | 0.774 | 0.609 | 0.731 | 0.600 | 0.624 | 0.646 | 0.553 |
| | Temporal | 0.892 | 0.905 | 0.925 | 0.814 | 0.886 | 0.842 | 0.881 | 0.903 | 0.655 |
| | Occipital | 0.907 | 0.921 | 0.907 | 0.780 | 0.850 | 0.747 | 0.838 | 0.865 | 0.739 |
| | Centro-parietal | 0.986 | 1.000 | 0.987 | 0.944 | 1.000 | 0.873 | 0.914 | 0.961 | 0.791 |
| Accuracy | Generalized | 0.860 | 0.877 | 0.877 | 0.737 | 0.772 | 0.772 | 0.789 | 0.789 | 0.719 |
| | Frontal | 0.739 | 0.739 | 0.783 | 0.609 | 0.739 | 0.652 | 0.630 | 0.674 | 0.565 |
| | Temporal | 0.943 | 0.957 | 0.971 | 0.843 | 0.943 | 0.800 | 0.900 | 0.929 | 0.529 |
| | Occipital | 0.829 | 0.854 | 0.829 | 0.780 | 0.829 | 0.829 | 0.756 | 0.780 | 0.829 |
| | Centro-parietal | 0.973 | 1.000 | 1.000 | 0.919 | 1.000 | 0.838 | 1.000 | 1.000 | 0.919 |
| Overall Accuracy | | 0.8725 | 0.8884 | 0.8964 | 0.7769 | 0.8566 | 0.7769 | 0.8167 | 0.8367 | 0.6853 |

**Description:** Reports class-specific precision, recall, F1 score, and accuracy for the five listed categories across the evaluated classifiers, together with overall accuracy.
**Abbreviations:** XGB, XGBoost; LGB, LightGBM; CatB, CatBoost; LR, logistic regression; LDA, linear discriminant analysis; DT, decision tree; ET, extra trees; RF, random forest; SVM, support vector machine.

**Table B.** Test-set classification performance for the EEG-only configuration.

| Performance Metrics | Seizure Onset Zone | XGB | LGB | CatB | LR | LDA | DT | ET | RF | SVM |
|---|---|---|---|---|---|---|---|---|---|---|
| Precision | Generalized | 0.943 | 0.961 | 0.981 | 0.827 | 0.944 | 0.906 | 0.940 | 0.941 | 0.683 |
| | Frontal | 0.800 | 0.776 | 0.833 | 0.600 | 0.784 | 0.569 | 0.723 | 0.736 | 0.591 |
| | Temporal | 0.882 | 0.883 | 0.893 | 0.787 | 0.914 | 0.881 | 0.789 | 0.861 | 0.717 |
| | Occipital | 1.000 | 0.972 | 0.974 | 0.739 | 0.975 | 0.761 | 0.973 | 0.973 | 0.558 |
| | Centro-parietal | 0.947 | 0.923 | 0.923 | 0.744 | 0.946 | 0.857 | 0.833 | 0.897 | 0.875 |
| Recall | Generalized | 0.862 | 0.845 | 0.879 | 0.741 | 0.879 | 0.828 | 0.810 | 0.828 | 0.741 |
| | Frontal | 0.870 | 0.826 | 0.870 | 0.652 | 0.870 | 0.630 | 0.739 | 0.848 | 0.565 |
| | Temporal | 0.957 | 0.971 | 0.957 | 0.686 | 0.914 | 0.843 | 0.857 | 0.886 | 0.543 |
| | Occipital | 0.833 | 0.833 | 0.881 | 0.810 | 0.929 | 0.833 | 0.857 | 0.857 | 0.690 |
| | Centro-parietal | 1.000 | 1.000 | 1.000 | 0.889 | 0.972 | 0.833 | 0.972 | 0.972 | 0.972 |
| F1 Score | Generalized | 0.901 | 0.899 | 0.927 | 0.782 | 0.911 | 0.865 | 0.870 | 0.881 | 0.711 |
| | Frontal | 0.833 | 0.800 | 0.851 | 0.625 | 0.825 | 0.598 | 0.731 | 0.788 | 0.578 |
| | Temporal | 0.918 | 0.925 | 0.924 | 0.733 | 0.914 | 0.861 | 0.822 | 0.873 | 0.618 |
| | Occipital | 0.909 | 0.897 | 0.925 | 0.773 | 0.951 | 0.795 | 0.911 | 0.911 | 0.617 |
| | Centro-parietal | 0.973 | 0.960 | 0.960 | 0.810 | 0.959 | 0.845 | 0.897 | 0.933 | 0.921 |
| Accuracy | Generalized | 0.862 | 0.845 | 0.879 | 0.741 | 0.879 | 0.828 | 0.810 | 0.828 | 0.741 |
| | Frontal | 0.870 | 0.826 | 0.870 | 0.652 | 0.870 | 0.630 | 0.739 | 0.848 | 0.565 |
| | Temporal | 0.957 | 0.971 | 0.957 | 0.686 | 0.914 | 0.843 | 0.857 | 0.886 | 0.543 |
| | Occipital | 0.833 | 0.833 | 0.881 | 0.810 | 0.929 | 0.833 | 0.857 | 0.857 | 0.690 |
| | Centro-parietal | 1.000 | 1.000 | 1.000 | 0.889 | 0.972 | 0.833 | 0.972 | 0.972 | 0.972 |
| **Overall Accuracy** | | 0.9048 | 0.8968 | 0.9167 | 0.7421 | 0.9087 | 0.7976 | 0.8413 | 0.8730 | 0.6786 |

**Description:** Reports class-specific precision, recall, F1 score, and accuracy for the five listed categories across the evaluated classifiers, together with overall accuracy.
**Abbreviations:** XGB, XGBoost; LGB, LightGBM; CatB, CatBoost; LR, logistic regression; LDA, linear discriminant analysis; DT, decision tree; ET, extra trees; RF, random forest; SVM, support vector machine.

**Table C.** Aggregated feature importance for the EEG-only configuration.

| Features | CatBoost | LGBM | XGB | DT | ET | RF | LR | LDA |
|---|---|---|---|---|---|---|---|---|
| **Power Beta** | 16.99 | 23.78 | 24.14 | 24.14 | 5.49 | 11.82 | ~0 | 2.13 |
| **Peak-to-Peak** | 9.38 | 12.02 | 12.29 | 12.29 | 10.90 | 12.65 | ~0 | 4.24 |
| **Power Alpha** | 5.78 | 4.72 | 5.38 | 5.38 | 4.52 | 6.18 | ~0 | 1.54 |
| **Hjorth Mobility** | 3.08 | 3.02 | 2.82 | 2.82 | 5.34 | 5.66 | ~0 | 3.46 |
| **Power Theta** | 7.28 | 12.28 | 12.29 | 12.29 | 6.08 | 10.39 | ~0 | 5.05 |
| **Waveform Length** | 6.77 | 2.99 | 3.83 | 3.83 | 9.11 | 7.93 | ~0 | 4.88 |
| **Approximate Entropy** | 4.67 | 5.70 | 4.96 | 4.96 | 8.94 | 6.38 | ~0 | 3.40 |
| **Standard Deviation** | 4.11 | 2.04 | 1.76 | 1.76 | 9.90 | 9.58 | ~0 | 9.89 |
| **Alpha-Beta Ratio** | 2.36 | 1.74 | 1.61 | 1.61 | 3.64 | 3.07 | 0.98 | 1.33 |
| **Skewness** | 4.50 | 5.45 | 5.44 | 5.44 | 1.31 | 1.58 | 0.30 | 0.52 |
| **Theta-Alpha Ratio** | 2.17 | 1.11 | 1.69 | 1.69 | 2.88 | 2.66 | 2.39 | 4.95 |
| **Permutation Entropy** | 3.71 | 3.35 | 4.31 | 4.31 | 4.77 | 2.65 | ~0 | 6.17 |
| **Kurtosis** | 7.32 | 4.84 | 4.72 | 4.72 | 1.95 | 1.99 | 3.93 | 0.93 |
| **Correlation Dimension** | 4.58 | 6.83 | 6.22 | 6.22 | 2.65 | 1.37 | 0.01 | 0.57 |
| **Sample Entropy** | 1.82 | 1.43 | 1.13 | 1.13 | 4.31 | 2.57 | 35.20 | 1.20 |
| **Power Delta** | 2.55 | 0.82 | 0.69 | 0.69 | 3.75 | 4.60 | ~0 | 2.63 |
| **Hjorth Complexity** | 2.60 | 2.17 | 1.03 | 1.03 | 1.21 | 1.29 | 0.41 | 1.65 |
| **Sample Shannon's Entropy** | 2.81 | 1.76 | 1.50 | 1.50 | 4.43 | 2.00 | 46.67 | 6.36 |
| **Median Frequency** | 2.22 | 1.37 | 1.53 | 1.53 | 1.30 | 0.78 | 1.03 | 0.67 |
| **Detrended Fluctuation Analysis** | 1.42 | 0.16 | 0.29 | 0.29 | 2.89 | 1.87 | 0.01 | 2.65 |
| **Hurst Exponent** | 0.74 | 0.28 | 0.55 | 0.55 | 0.64 | 0.41 | ~0 | 0.79 |
| **Spectral Entropy** | 0.68 | 0.25 | 0.15 | 0.15 | 1.89 | 0.81 | 0.22 | 1.27 |
| **Delta-Theta Ratio** | 0.30 | 0.39 | 0.58 | 0.58 | 0.83 | 0.84 | 7.51 | 1.10 |
| **Peak Frequency** | 0.64 | 0.74 | 0.61 | 0.61 | 0.48 | 0.25 | 1.25 | 0.27 |
| **Median** | 1.16 | 0.59 | 0.27 | 0.27 | 0.52 | 0.40 | ~0 | 0.35 |
| **Mean** | 0.25 | 0.05 | 0.08 | 0.08 | 0.14 | 0.17 | ~0 | 31.87 |

**Description:** Reports normalized SHAP importance percentages aggregated by feature family across the evaluated classifiers. **Abbreviations:** SHAP, SHapley Additive exPlanations; LGBM, LightGBM; XGB, XGBoost; DT, decision tree; ET, extra trees; RF, random forest; LR, logistic regression; LDA, linear discriminant analysis.

**Table D.** Channel-wise importance for the EEG-only configuration.

| Lead | Indication | CatBoost | LGBM | XGB | DT | ET | RF | LR | LDA |
|---|---|---|---|---|---|---|---|---|---|
| **Lead 1** | EEG Fp1 | 7.47 | 9.93 | 10.29 | 10.29 | 5.28 | 7.25 | 5.00 | 4.77 |
| **Lead 2** | EEG F3 | 8.65 | 6.41 | 6.20 | 6.20 | 4.47 | 5.73 | 6.33 | 4.08 |
| **Lead 3** | EEG C3 | 3.47 | 1.80 | 2.13 | 2.13 | 6.07 | 5.90 | 5.72 | 4.20 |
| **Lead 4** | EEG P3 | 5.38 | 6.98 | 8.03 | 8.03 | 7.44 | 7.88 | 7.15 | 4.34 |
| **Lead 5** | EEG O1 | 0.81 | 0.31 | 0.29 | 0.29 | 2.58 | 1.13 | 2.74 | 3.48 |
| **Lead 6** | EEG F7 | 3.14 | 2.27 | 2.26 | 2.26 | 3.87 | 4.03 | 2.62 | 3.40 |
| **Lead 7** | EEG T3 | 2.15 | 3.18 | 3.63 | 3.63 | 4.34 | 2.97 | 3.99 | 3.12 |
| **Lead 8** | EEG T5 | 2.77 | 3.68 | 4.16 | 4.16 | 4.19 | 3.53 | 4.95 | 4.17 |
| **Lead 9** | EEG Fz | 10.66 | 11.16 | 11.58 | 11.58 | 8.56 | 11.16 | 5.99 | 4.98 |
| **Lead 10** | EEG Cz | 3.97 | 3.66 | 2.66 | 2.66 | 5.84 | 4.93 | 4.91 | 4.90 |
| **Lead 11** | EEG Pz | 1.14 | 1.95 | 1.68 | 1.68 | 1.94 | 1.42 | 3.82 | 3.17 |
| **Lead 12** | EEG Fp2 | 2.72 | 1.58 | 1.64 | 1.64 | 2.98 | 1.98 | 2.15 | 3.61 |
| **Lead 13** | EEG F4 | 1.59 | 0.53 | 0.52 | 0.52 | 1.95 | 1.19 | 4.68 | 2.98 |
| **Lead 14** | EEG C4 | 2.99 | 2.37 | 2.37 | 2.37 | 3.13 | 3.37 | 3.65 | 3.05 |
| **Lead 15** | EEG P4 | 3.48 | 3.00 | 3.01 | 3.01 | 6.09 | 6.80 | 3.34 | 3.92 |
| **Lead 16** | EEG O2 | 2.40 | 1.72 | 1.88 | 1.88 | 2.59 | 1.61 | 3.50 | 4.43 |
| **Lead 17** | EEG F8 | 6.19 | 8.09 | 8.41 | 8.41 | 8.73 | 9.38 | 4.97 | 4.23 |
| **Lead 18** | EEG T4 | 3.15 | 2.26 | 2.34 | 2.34 | 4.44 | 3.25 | 5.28 | 3.40 |
| **Lead 19** | EEG T6 | 4.21 | 5.80 | 5.95 | 5.95 | 4.48 | 3.42 | 2.68 | 3.96 |
| **Lead 26** | EMG (Left deltoid 1) | 5.89 | 7.13 | 7.02 | 7.02 | 3.11 | 4.31 | 4.89 | 5.42 |
| **Lead 27** | EMG (Left deltoid 2) | 6.61 | 4.97 | 3.77 | 3.77 | 3.04 | 4.03 | 6.10 | 5.01 |
| **Lead 28** | EMG (Right deltoid 1) | 4.18 | 5.47 | 4.87 | 4.87 | 2.45 | 2.26 | 2.75 | 10.44 |
| **Lead 29** | EMG (Right deltoid 2) | 6.89 | 5.62 | 5.20 | 5.20 | 2.31 | 2.36 | 2.67 | 4.81 |

**Description:** Reports normalized SHAP importance percentages aggregated by recording channel across the evaluated classifiers. **Abbreviations:** SHAP, SHapley Additive exPlanations; LGBM, LightGBM; XGB, XGBoost; DT, decision tree; ET, extra trees; RF, random forest; LR, logistic regression; LDA, linear discriminant analysis.

**Table E.** Top channel-feature SHAP attributions for CatBoost.

| Rank | Feature | SHAP Importance (%) |
|---|---|---|
| **1** | lead9_power_theta | 2.31 |
| **2** | lead2_power_beta | 2.21 |
| **3** | lead19_power_beta | 2.16 |
| **4** | lead1_power_beta | 1.94 |
| **5** | lead9_waveform_length | 1.93 |
| **6** | lead17_power_beta | 1.68 |
| **7** | lead9_ptp | 1.66 |
| **8** | lead4_kurtosis | 1.35 |
| **9** | lead1_power_alpha | 1.27 |
| **10** | lead9_power_alpha | 1.23 |

**Description:** Ranks the ten channel-feature variables with the largest normalized SHAP importance values for the CatBoost model. **Abbreviations:** SHAP, SHapley Additive exPlanations.

**Table F.** Top channel-feature SHAP attributions for LightGBM.

| Rank | Feature | SHAP Importance (%) |
|---|---|---|
| **1** | lead9_power_theta | 4.72 |
| **2** | lead1_power_beta | 3.99 |
| **3** | lead26_power_beta | 2.80 |
| **4** | lead9_power_alpha | 2.63 |
| **5** | lead2_power_beta | 2.59 |
| **6** | lead19_power_beta | 2.06 |
| **7** | lead17_approx_entropy | 1.90 |
| **8** | lead4_power_theta | 1.88 |
| **9** | lead8_power_beta | 1.81 |
| **10** | lead17_power_beta | 1.79 |

**Description:** Ranks the ten channel-feature variables with the largest normalized SHAP importance values for the LightGBM model. **Abbreviations:** SHAP, SHapley Additive exPlanations.

**Table G.** Top channel-feature SHAP attributions for XGBoost.

| Rank | Feature | SHAP Importance (%) |
|---|---|---|
| **1** | lead9_power_theta | 4.99 |
| **2** | lead1_power_beta | 4.80 |
| **3** | lead9_power_alpha | 2.79 |
| **4** | lead19_power_beta | 2.37 |
| **5** | lead26_power_beta | 2.21 |
| **6** | lead2_power_beta | 2.17 |
| **7** | lead17_approx_entropy | 1.92 |
| **8** | lead4_power_theta | 1.89 |
| **9** | lead8_power_beta | 1.87 |
| **10** | lead17_ptp | 1.86 |

**Description:** Ranks the ten channel-feature variables with the largest normalized SHAP importance values for the XGBoost model. **Abbreviations:** SHAP, SHapley Additive exPlanations.

**Table H.** Top channel-feature SHAP attributions for the decision tree.

| Rank | Feature | SHAP Importance (%) |
|---|---|---|
| **1** | lead9_power_theta | 4.99 |
| **2** | lead1_power_beta | 4.80 |
| **3** | lead9_power_alpha | 2.79 |
| **4** | lead19_power_beta | 2.37 |
| **5** | lead26_power_beta | 2.21 |
| **6** | lead2_power_beta | 2.17 |
| **7** | lead17_approx_entropy | 1.92 |
| **8** | lead4_power_theta | 1.89 |
| **9** | lead8_power_beta | 1.87 |
| **10** | lead17_ptp | 1.86 |

**Description:** Ranks the ten channel-feature variables with the largest normalized SHAP importance values for the decision-tree model. **Abbreviations:** SHAP, SHapley Additive exPlanations.

**Table I.** Top channel-feature SHAP attributions for extra trees.

| Rank | Feature | SHAP Importance (%) |
|---|---|---|
| **1** | lead9_ptp | 1.17 |
| **2** | lead9_waveform_length | 1.14 |
| **3** | lead17_std | 1.09 |
| **4** | lead9_std | 0.99 |
| **5** | lead15_std | 0.98 |
| **6** | lead3_ptp | 0.96 |
| **7** | lead15_power_theta | 0.94 |
| **8** | lead17_ptp | 0.91 |
| **9** | lead10_waveform_length | 0.86 |
| **10** | lead4_std | 0.86 |

**Description:** Ranks the ten channel-feature variables with the largest normalized SHAP importance values for the extra-trees model. **Abbreviations:** SHAP, SHapley Additive exPlanations.

**Table J.** Top channel-feature SHAP attributions for random forest.

| Rank | Feature | SHAP Importance (%) |
|---|---|---|
| 1 | lead9_power_theta | 2.34 |
| 2 | lead9_ptp | 1.88 |
| 3 | lead15_power_theta | 1.83 |
| 4 | lead9_std | 1.52 |
| 5 | lead15_ptp | 1.50 |
| 6 | lead1_power_beta | 1.47 |
| 7 | lead9_power_alpha | 1.38 |
| 8 | lead2_power_beta | 1.35 |
| 9 | lead17_std | 1.32 |
| 10 | lead17_approx_entropy | 1.24 |

**Description:** Ranks the ten channel-feature variables with the largest normalized SHAP importance values for the random-forest model. **Abbreviations:** SHAP, SHapley Additive exPlanations.

**Table K.** Top channel-feature SHAP attributions for logistic regression.

| Rank | Feature | SHAP Importance (%) |
|---|---|---|
| **1** | lead4_ssc | 3.96 |
| **2** | lead2_ssc | 3.79 |
| **3** | lead3_ssc | 3.42 |
| **4** | lead9_ssc | 2.89 |
| **5** | lead27_ssc | 2.81 |
| **6** | lead8_sample_entropy | 2.68 |
| **7** | lead18_ssc | 2.48 |
| **8** | lead1_ssc | 2.30 |
| **9** | lead17_ssc | 2.28 |
| **10** | lead4_sample_entropy | 2.17 |

**Description:** Ranks the ten channel-feature variables with the largest normalized SHAP importance values for the logistic-regression model. **Abbreviations:** SHAP, SHapley Additive exPlanations.

**Table L.** Top channel-feature SHAP attributions for linear discriminant analysis.

| Rank | Feature | SHAP Importance (%) |
|---|---|---|
| **1** | lead28_theta_alpha_ratio | 3.52 |
| **2** | lead28_power_theta | 3.44 |
| **3** | lead2_mean | 2.44 |
| **4** | lead1_mean | 2.44 |
| **5** | lead10_mean | 2.08 |
| **6** | lead17_mean | 2.01 |
| **7** | lead9_mean | 2.01 |
| **8** | lead4_mean | 1.89 |
| **9** | lead16_mean | 1.82 |
| **10** | lead3_mean | 1.72 |

**Description:** Ranks the ten channel-feature variables with the largest normalized SHAP importance values for the linear-discriminant-analysis model. **Abbreviations:** SHAP, SHapley Additive exPlanations.

### S3 Table

#### EEG+ECG validation/test performance and SHAP importance results

This supporting file contains validation- and test-set performance metrics, aggregated feature-family and channel-wise SHAP importance, and model-specific top-ten channel-feature SHAP rankings for the EEG+ECG configuration. Tables A-L are presented as editable Word tables.

##### Contents and table descriptions

| Internal table | Description |
|---|---|
| Table A | Validation-set classification performance for the EEG+ECG configuration. Reports class-specific precision, recall, F1 score, and accuracy for the five listed categories across the evaluated classifiers, together with overall accuracy. |
| Table B | Test-set classification performance for the EEG+ECG configuration. Reports class-specific precision, recall, F1 score, and accuracy for the five listed categories across the evaluated classifiers, together with overall accuracy. |
| Table C | Aggregated feature importance for the EEG+ECG configuration. Reports normalized SHAP importance percentages aggregated by feature family across the evaluated classifiers. |
| Table D | Channel-wise importance for the EEG+ECG configuration. Reports normalized SHAP importance percentages aggregated by recording channel across the evaluated classifiers. |
| Table E | Top channel-feature SHAP attributions for CatBoost. Ranks the ten channel-feature variables with the largest normalized SHAP importance values for the CatBoost model. |
| Table F | Top channel-feature SHAP attributions for LightGBM. Ranks the ten channel-feature variables with the largest normalized SHAP importance values for the LightGBM model. |
| Table G | Top channel-feature SHAP attributions for XGBoost. Ranks the ten channel-feature variables with the largest normalized SHAP importance values for the XGBoost model. |
| Table H | Top channel-feature SHAP attributions for the decision tree. Ranks the ten channel-feature variables with the largest normalized SHAP importance values for the decision-tree model. |
| Table I | Top channel-feature SHAP attributions for extra trees. Ranks the ten channel-feature variables with the largest normalized SHAP importance values for the extra-trees model. |
| Table J | Top channel-feature SHAP attributions for random forest. Ranks the ten channel-feature variables with the largest normalized SHAP importance values for the random-forest model. |
| Table K | Top channel-feature SHAP attributions for logistic regression. Ranks the ten channel-feature variables with the largest normalized SHAP importance values for the logistic-regression model. |
| Table L | Top channel-feature SHAP attributions for linear discriminant analysis. Ranks the ten channel-feature variables with the largest normalized SHAP importance values for the linear-discriminant-analysis model. |

**Formatting note:** All tables remain editable, cell-based objects. Numerical values and source table text were retained; only layout, captions, and descriptive legends were added.

**Table A.** Validation-set classification performance for the EEG+ECG configuration.

| Performance Metrics | Seizure Onset Zone | CatB | LGB | XGB | DT | ET | LR | LDA | SVM |
|---|---|---|---|---|---|---|---|---|---|
| Precision | Generalized | 0.907 | 0.862 | 0.893 | 0.870 | 0.833 | 0.865 | 0.902 | 0.613 |
| | Frontal | 0.818 | 0.756 | 0.795 | 0.617 | 0.640 | 0.696 | 0.744 | 0.431 |
| | temporal | 0.850 | 0.873 | 0.896 | 0.808 | 0.877 | 0.763 | 0.821 | 0.811 |
| | occipital | 1.000 | 1.000 | 1.000 | 0.829 | 0.902 | 0.889 | 1.000 | 0.810 |
| | Centro-parietal | 0.972 | 1.000 | 1.000 | 0.889 | 0.970 | 0.780 | 0.972 | 0.767 |
| Recall | Generalized | 0.860 | 0.877 | 0.877 | 0.825 | 0.789 | 0.789 | 0.807 | 0.667 |
| | Frontal | 0.783 | 0.674 | 0.761 | 0.630 | 0.696 | 0.696 | 0.696 | 0.478 |
| | temporal | 0.971 | 0.986 | 0.986 | 0.843 | 0.914 | 0.829 | 0.986 | 0.614 |
| | occipital | 1.000 | 1.000 | 1.000 | 0.919 | 1.000 | 0.865 | 1.000 | 0.919 |
| | Centro-parietal | 0.854 | 0.878 | 0.902 | 0.780 | 0.780 | 0.780 | 0.854 | 0.805 |
| F1 Score | Generalized | 0.883 | 0.870 | 0.885 | 0.847 | 0.811 | 0.826 | 0.852 | 0.639 |
| | Frontal | 0.800 | 0.713 | 0.778 | 0.624 | 0.667 | 0.696 | 0.719 | 0.454 |
| | temporal | 0.907 | 0.926 | 0.939 | 0.825 | 0.895 | 0.795 | 0.896 | 0.699 |
| | occipital | 1.000 | 1.000 | 1.000 | 0.872 | 0.949 | 0.877 | 1.000 | 0.861 |
| | Centro-parietal | 0.909 | 0.935 | 0.949 | 0.831 | 0.865 | 0.780 | 0.909 | 0.786 |
| Accuracy | Generalized | 0.860 | 0.877 | 0.877 | 0.825 | 0.789 | 0.789 | 0.807 | 0.667 |
| | Frontal | 0.783 | 0.674 | 0.761 | 0.630 | 0.696 | 0.696 | 0.696 | 0.478 |
| | temporal | 0.971 | 0.986 | 0.986 | 0.843 | 0.914 | 0.829 | 0.986 | 0.614 |
| | occipital | 1.000 | 1.000 | 1.000 | 0.919 | 1.000 | 0.865 | 1.000 | 0.919 |
| | Centro-parietal | 0.854 | 0.878 | 0.902 | 0.780 | 0.780 | 0.780 | 0.854 | 0.805 |
| **Overall Accuracy** | | 0.896 | 0.888 | 0.908 | 0.801 | 0.837 | 0.793 | 0.873 | 0.677 |

**Description:** Reports class-specific precision, recall, F1 score, and accuracy for the five listed categories across the evaluated classifiers, together with overall accuracy.
**Abbreviations:** CatB, CatBoost; LGB, LightGBM; XGB, XGBoost; DT, decision tree; ET, extra trees; LR, logistic regression; LDA, linear discriminant analysis; SVM, support vector machine.

**Table B.** Test-set classification performance for the EEG+ECG configuration.

| Performance Metrics | Seizure Onset Zone | CatB | LGB | XGB | DT | ET | LR | LDA | SVM |
|---|---|---|---|---|---|---|---|---|---|
| Precision | Generalized | 0.947 | 0.907 | 0.923 | 0.882 | 0.922 | 0.780 | 0.959 | 0.667 |
| | Frontal | 0.894 | 0.787 | 0.776 | 0.564 | 0.714 | 0.638 | 0.804 | 0.396 |
| | temporal | 0.942 | 0.877 | 0.903 | 0.857 | 0.881 | 0.843 | 0.882 | 0.692 |
| | occipital | 0.947 | 0.919 | 0.921 | 0.838 | 0.897 | 0.892 | 0.973 | 0.800 |
| | Centro-parietal | 0.927 | 0.927 | 0.927 | 0.872 | 0.949 | 0.821 | 0.974 | 0.660 |
| Recall | Generalized | 0.931 | 0.845 | 0.828 | 0.776 | 0.810 | 0.793 | 0.810 | 0.690 |
| | Frontal | 0.913 | 0.804 | 0.826 | 0.674 | 0.870 | 0.652 | 0.891 | 0.457 |
| | temporal | 0.929 | 0.914 | 0.929 | 0.857 | 0.843 | 0.843 | 0.957 | 0.514 |
| | occipital | 1.000 | 0.944 | 0.972 | 0.861 | 0.972 | 0.917 | 1.000 | 0.889 |
| | Centro-parietal | 0.905 | 0.905 | 0.905 | 0.810 | 0.881 | 0.762 | 0.905 | 0.738 |
| F1 Score | Generalized | 0.939 | 0.875 | 0.873 | 0.826 | 0.862 | 0.786 | 0.879 | 0.678 |
| | Frontal | 0.903 | 0.796 | 0.800 | 0.614 | 0.784 | 0.645 | 0.845 | 0.424 |
| | temporal | 0.935 | 0.895 | 0.915 | 0.857 | 0.861 | 0.843 | 0.918 | 0.590 |
| | occipital | 0.973 | 0.932 | 0.946 | 0.849 | 0.933 | 0.904 | 0.986 | 0.842 |
| | Centro-parietal | 0.916 | 0.916 | 0.916 | 0.840 | 0.914 | 0.790 | 0.938 | 0.697 |
| Accuracy | Generalized | 0.931 | 0.845 | 0.828 | 0.776 | 0.810 | 0.793 | 0.810 | 0.690 |
| | Frontal | 0.913 | 0.804 | 0.826 | 0.674 | 0.870 | 0.652 | 0.891 | 0.457 |
| | temporal | 0.929 | 0.914 | 0.929 | 0.857 | 0.843 | 0.843 | 0.957 | 0.514 |
| | occipital | 1.000 | 0.944 | 0.972 | 0.861 | 0.972 | 0.917 | 1.000 | 0.889 |
| | Centro-parietal | 0.905 | 0.905 | 0.905 | 0.810 | 0.881 | 0.762 | 0.905 | 0.738 |
| **Overall Accuracy** | | 0.933 | 0.881 | 0.889 | 0.798 | 0.865 | 0.794 | 0.909 | 0.635 |

**Description:** Reports class-specific precision, recall, F1 score, and accuracy for the five listed categories across the evaluated classifiers, together with overall accuracy.
**Abbreviations:** CatB, CatBoost; LGB, LightGBM; XGB, XGBoost; DT, decision tree; ET, extra trees; LR, logistic regression; LDA, linear discriminant analysis; SVM, support vector machine.

**Table C.** Aggregated feature importance for the EEG+ECG configuration.

| Features | CatBoost | LGBM | XGB | DT | ET | RF | LR | LDA |
|---|---|---|---|---|---|---|---|---|
| **Power Beta** | 18.76 | 22.72 | 23.31 | 23.31 | 5.30 | 11.87 | ~0 | 2.16 |
| **Peak-to-Peak** | 9.72 | 8.95 | 8.55 | 8.55 | 10.29 | 11.48 | ~0 | 2.74 |
| **Power Alpha** | 7.28 | 7.36 | 7.45 | 7.45 | 5.20 | 6.52 | ~0 | 1.65 |
| **Hjorth Mobility** | 3.47 | 4.53 | 4.34 | 4.34 | 5.48 | 6.08 | ~0 | 4.82 |
| **Power Theta** | 8.08 | 9.50 | 9.77 | 9.77 | 7.08 | 11.83 | ~0 | 2.06 |
| **Waveform Length** | 4.15 | 2.09 | 3.03 | 3.03 | 7.69 | 6.55 | ~0 | 5.35 |
| **Approximate Entropy** | 3.36 | 3.62 | 3.58 | 3.58 | 7.49 | 5.99 | 0.01 | 4.23 |
| **Standard Deviation** | 2.11 | 2.14 | 2.72 | 2.72 | 9.43 | 7.20 | ~0 | 8.52 |
| **Alpha-Beta Ratio** | 7.98 | 5.72 | 5.12 | 5.12 | 4.48 | 4.21 | 1.55 | 1.40 |
| **Skewness** | 4.57 | 4.23 | 4.19 | 4.19 | 1.88 | 2.44 | 0.74 | 0.71 |
| **Theta-Alpha Ratio** | 5.69 | 3.54 | 3.46 | 3.46 | 2.86 | 2.63 | 3.95 | 1.40 |
| **Permutation Entropy** | 3.91 | 3.46 | 4.56 | 4.56 | 4.34 | 2.53 | 0.01 | 8.39 |
| **Kurtosis** | 1.93 | 2.36 | 2.04 | 2.04 | 1.46 | 1.24 | 6.68 | 1.04 |
| **Correlation Dimension** | 3.64 | 6.31 | 5.31 | 5.31 | 2.73 | 1.66 | 0.04 | 0.68 |
| **Sample Entropy** | 2.43 | 1.82 | 1.24 | 1.24 | 4.03 | 2.81 | 28.96 | 1.57 |
| **Power Delta** | 1.67 | 0.17 | 0.52 | 0.52 | 4.07 | 4.61 | ~0 | 2.89 |
| **Hjorth Complexity** | 1.08 | 0.63 | 0.61 | 0.61 | 1.83 | 1.47 | 0.90 | 2.47 |
| **Slope Sign Changes (SSC)** | 1.01 | 2.65 | 2.60 | 2.60 | 4.12 | 2.06 | 39.46 | 8.45 |
| **Median Frequency** | 1.61 | 1.35 | 1.23 | 1.23 | 1.49 | 0.68 | 2.62 | 1.11 |
| **Detrended Fluctuation Analysis** | 0.90 | 0.56 | 0.47 | 0.47 | 3.50 | 2.13 | 0.05 | 3.34 |
| **Hurst Exponent** | 0.11 | 0.63 | 0.43 | 0.43 | 0.70 | 0.45 | 0.01 | 0.75 |
| **Spectral Entropy** | 1.04 | 0.77 | 0.54 | 0.54 | 1.95 | 0.90 | 0.67 | 1.63 |
| **Delta-Theta Ratio** | 2.74 | 2.36 | 2.40 | 2.40 | 1.13 | 1.04 | 12.59 | 0.91 |
| **Peak Frequency** | 1.16 | 1.61 | 1.58 | 1.58 | 0.59 | 0.65 | 1.76 | 0.34 |
| **Median** | 1.32 | 0.92 | 0.89 | 0.89 | 0.75 | 0.80 | ~0 | 0.41 |
| **Mean** | 0.26 | 0.00 | 0.04 | 0.04 | 0.13 | 0.16 | ~0 | 30.99 |

**Description:** Reports normalized SHAP importance percentages aggregated by feature family across the evaluated classifiers. **Abbreviations:** SHAP, SHapley Additive exPlanations; LGBM, LightGBM; XGB, XGBoost; DT, decision tree; ET, extra trees; RF, random forest; LR, logistic regression; LDA, linear discriminant analysis.

**Table D.** Channel-wise importance for the EEG+ECG configuration.

| Lead | Indication | CatBoost | LGBM | XGB | DT | ET | RF | LR | LDA |
|---|---|---|---|---|---|---|---|---|---|
| **Lead 1** | EEG Fp1 | 7.32 | 9.51 | 9.12 | 9.12 | 5.76 | 5.91 | 6.52 | 5.74 |
| **Lead 2** | EEG F3 | 5.80 | 4.43 | 4.02 | 4.02 | 4.96 | 5.96 | 6.58 | 4.71 |
| **Lead 3** | EEG C3 | 3.45 | 2.17 | 2.29 | 2.29 | 5.81 | 6.35 | 5.19 | 5.29 |
| **Lead 4** | EEG P3 | 6.65 | 6.77 | 7.81 | 7.81 | 6.98 | 7.06 | 7.97 | 5.10 |
| **Lead 5** | EEG O1 | 0.46 | 0.09 | 0.36 | 0.36 | 2.54 | 1.07 | 2.36 | 4.36 |
| **Lead 6** | EEG F7 | 1.88 | 2.29 | 2.42 | 2.42 | 3.98 | 3.89 | 3.51 | 3.97 |
| **Lead 7** | EEG T3 | 1.56 | 2.73 | 2.69 | 2.69 | 3.91 | 2.47 | 3.87 | 4.13 |
| **Lead 8** | EEG T5 | 4.10 | 3.55 | 3.05 | 3.05 | 4.24 | 3.40 | 4.51 | 5.64 |
| **Lead 9** | EEG Fz | 8.24 | 7.54 | 7.89 | 7.89 | 7.78 | 11.31 | 6.12 | 6.10 |
| **Lead 10** | EEG Cz | 3.44 | 2.32 | 1.54 | 1.54 | 5.63 | 4.58 | 5.53 | 5.92 |
| **Lead 11** | EEG Pz | 1.93 | 1.62 | 1.72 | 1.72 | 2.17 | 1.55 | 3.87 | 3.82 |
| **Lead 12** | EEG Fp2 | 1.96 | 1.11 | 1.18 | 1.18 | 2.64 | 1.74 | 2.76 | 4.69 |
| **Lead 13** | EEG F4 | 0.91 | 0.03 | 0.19 | 0.19 | 1.93 | 1.20 | 3.58 | 3.96 |
| **Lead 14** | EEG C4 | 1.72 | 1.86 | 1.54 | 1.54 | 3.03 | 2.65 | 2.95 | 3.19 |
| **Lead 15** | EEG P4 | 4.57 | 3.13 | 2.97 | 2.97 | 6.07 | 7.23 | 3.78 | 5.16 |
| **Lead 16** | EEG O2 | 1.96 | 1.29 | 1.34 | 1.34 | 2.70 | 1.55 | 2.06 | 5.09 |
| **Lead 17** | EEG F8 | 7.03 | 6.14 | 6.81 | 6.81 | 8.18 | 9.41 | 5.14 | 5.13 |
| **Lead 18** | EEG T4 | 3.72 | 1.33 | 1.73 | 1.73 | 3.54 | 2.82 | 6.73 | 3.62 |
| **Lead 19** | EEG T6 | 2.39 | 3.40 | 4.30 | 4.30 | 4.13 | 3.08 | 3.27 | 5.01 |
| **Lead 24** | ECG channel 1 | 15.79 | 17.89 | 16.07 | 16.07 | 7.02 | 7.40 | 6.36 | 5.54 |
| **Lead 25** | ECG channel 2 | 15.12 | 20.80 | 20.95 | 20.95 | 7.01 | 9.37 | 7.35 | 3.83 |

**Description:** Reports normalized SHAP importance percentages aggregated by recording channel across the evaluated classifiers. **Abbreviations:** SHAP, SHapley Additive exPlanations; LGBM, LightGBM; XGB, XGBoost; DT, decision tree; ET, extra trees; RF, random forest; LR, logistic regression; LDA, linear discriminant analysis.

**Table E.** Top channel-feature SHAP attributions for CatBoost.

| Rank | Feature | SHAP Importance (%) |
|---|---|---|
| **1** | lead24_alpha_beta_ratio | 3.79 |
| **2** | lead1_power_beta | 3.33 |
| **3** | lead25_power_alpha | 2.40 |
| **4** | lead25_theta_alpha_ratio | 2.31 |
| **5** | lead9_ptp | 2.07 |
| **6** | lead9_power_theta | 2.05 |
| **7** | lead25_power_beta | 2.03 |
| **8** | lead25_alpha_beta_ratio | 1.86 |
| **9** | lead8_power_beta | 1.85 |
| **10** | lead2_power_beta | 1.53 |

**Description:** Ranks the ten channel-feature variables with the largest normalized SHAP importance values for the CatBoost model. **Abbreviations:** SHAP, SHapley Additive exPlanations.

**Table F.** Top channel-feature SHAP attributions for LightGBM.

| Rank | Feature | SHAP Importance (%) |
|---|---|---|
| **1** | lead25_power_beta | 5.68 |
| **2** | lead1_power_beta | 4.97 |
| **3** | lead25_power_alpha | 4.13 |
| **4** | lead24_alpha_beta_ratio | 3.18 |
| **5** | lead25_perm_entropy | 2.42 |
| **6** | lead9_power_theta | 2.40 |
| **7** | lead25_theta_alpha_ratio | 2.19 |
| **8** | lead25_ssc | 2.07 |
| **9** | lead24_hjorth_mobility | 1.93 |
| **10** | lead9_power_alpha | 1.82 |

**Description:** Ranks the ten channel-feature variables with the largest normalized SHAP importance values for the LightGBM model. **Abbreviations:** SHAP, SHapley Additive exPlanations.

**Table G.** Top channel-feature SHAP attributions for XGBoost.

| Rank | Feature | SHAP Importance (%) |
|---|---|---|
| 1 | lead25_power_beta | 6.10 |
| 2 | lead1_power_beta | 4.62 |
| 3 | lead25_power_alpha | 4.22 |
| 4 | lead9_power_theta | 3.11 |
| 5 | lead25_perm_entropy | 2.83 |
| 6 | lead24_alpha_beta_ratio | 2.80 |
| 7 | lead17_approx_entropy | 2.19 |
| 8 | lead4_power_theta | 2.00 |
| 9 | lead25_ssc | 1.81 |
| 10 | lead25_theta_alpha_ratio | 1.79 |

**Description:** Ranks the ten channel-feature variables with the largest normalized SHAP importance values for the XGBoost model. **Abbreviations:** SHAP, SHapley Additive exPlanations.

**Table H.** Top channel-feature SHAP attributions for the decision tree.

| Rank | Feature | SHAP Importance (%) |
|---|---|---|
| **1** | lead25_power_beta | 6.10 |
| **2** | lead1_power_beta | 4.62 |
| **3** | lead25_power_alpha | 4.22 |
| **4** | lead9_power_theta | 3.11 |
| **5** | lead25_perm_entropy | 2.83 |
| **6** | lead24_alpha_beta_ratio | 2.80 |
| **7** | lead17_approx_entropy | 2.19 |
| **8** | lead4_power_theta | 2.00 |
| **9** | lead25_ssc | 1.81 |
| **10** | lead25_theta_alpha_ratio | 1.79 |

**Description:** Ranks the ten channel-feature variables with the largest normalized SHAP importance values for the decision-tree model. **Abbreviations:** SHAP, SHapley Additive exPlanations.

**Table I.** Top channel-feature SHAP attributions for extra trees.

| Rank | Feature | SHAP Importance (%) |
|---|---|---|
| **1** | lead9_ptp | 1.37 |
| **2** | lead17_std | 1.28 |
| **3** | lead15_ptp | 1.00 |
| **4** | lead24_alpha_beta_ratio | 0.99 |
| **5** | lead4_std | 0.92 |
| **6** | lead3_ptp | 0.92 |
| **7** | lead9_std | 0.90 |
| **8** | lead25_power_alpha | 0.85 |
| **9** | lead4_approx_entropy | 0.84 |
| **10** | lead10_std | 0.82 |

**Description:** Ranks the ten channel-feature variables with the largest normalized SHAP importance values for the extra-trees model. **Abbreviations:** SHAP, SHapley Additive exPlanations.

**Table J.** Top channel-feature SHAP attributions for random forest.

| Rank | Feature | SHAP Importance (%) |
|---|---|---|
| **1** | lead9_power_theta | 2.46 |
| **2** | lead15_power_theta | 2.21 |
| **3** | lead9_ptp | 2.07 |
| **4** | lead17_approx_entropy | 1.69 |
| **5** | lead10_power_theta | 1.55 |
| **6** | lead15_ptp | 1.53 |
| **7** | lead17_ptp | 1.41 |
| **8** | lead25_power_beta | 1.35 |
| **9** | lead9_power_alpha | 1.35 |
| **10** | lead24_alpha_beta_ratio | 1.32 |

**Description:** Ranks the ten channel-feature variables with the largest normalized SHAP importance values for the random-forest model. **Abbreviations:** SHAP, SHapley Additive exPlanations.

**Table K.** Top channel-feature SHAP attributions for logistic regression.

| Rank | Feature | SHAP Importance (%) |
|---|---|---|
| **1** | lead4_ssc | 4.19 |
| **2** | lead2_ssc | 3.62 |
| **3** | lead18_sample_entropy | 2.89 |
| **4** | lead9_ssc | 2.78 |
| **5** | lead3_ssc | 2.64 |
| **6** | lead1_ssc | 2.64 |
| **7** | lead10_ssc | 2.45 |
| **8** | lead25_ssc | 2.37 |
| **9** | lead17_ssc | 2.33 |
| **10** | lead8_ssc | 2.13 |

**Description:** Ranks the ten channel-feature variables with the largest normalized SHAP importance values for the logistic-regression model. **Abbreviations:** SHAP, SHapley Additive exPlanations.

**Table L.** Top channel-feature SHAP attributions for linear discriminant analysis.

| Rank | Feature | SHAP Importance (%) |
|---|---|---|
| **1** | lead2_mean | 2.38 |
| **2** | lead1_mean | 2.36 |
| **3** | lead10_mean | 2.05 |
| **4** | lead17_mean | 1.94 |
| **5** | lead9_mean | 1.92 |
| **6** | lead4_mean | 1.89 |
| **7** | lead16_mean | 1.78 |
| **8** | lead3_mean | 1.68 |
| **9** | lead12_mean | 1.57 |
| **10** | lead11_mean | 1.57 |

**Description:** Ranks the ten channel-feature variables with the largest normalized SHAP importance values for the linear-discriminant-analysis model. **Abbreviations:** SHAP, SHapley Additive exPlanations.
